# Rsc1 DNA-binding bromodomain drives RSC activity at A-rich promoters

**DOI:** 10.1101/2025.03.24.644971

**Authors:** Camille Sayou, Oriane Viravaux, Jordan Lyonnet, Jérémy Lucas, Raphaël Dupeyron, Emmanuel Thévenon, Julie Marais, Annie Adrait, Yohann Couté, François Parcy, Carlo Petosa, Jérôme Govin

## Abstract

In yeast, the SWI/SNF chromatin remodeler RSC orchestrates nucleosome positioning at promoters, yet the molecular determinants guiding RSC to chromatin remain unclear. Here, we investigate the RSC subunit Rsc1 in *Candida albicans* and *Saccharomyces cerevisiae*. We identify a non-canonical role for its second bromodomain, Rsc1-BD2, which lacks a functional acetyllysine-binding pocket. Instead, Rsc1-BD2, together with the adjacent AT-hook and BAH domains, forms a conserved DNA-binding surface that is critical for RSC-dependent nucleosome remodeling and essential for viability in both species. Genome-wide motif analysis reveals that promoter regions are enriched in polyA sequences, whose distribution correlates with nucleosome positioning and gene expression. We show that Rsc1 preferentially binds polyA motifs and that nucleosome positioning at polyA-rich promoters depends on Rsc1 DNA binding module. These findings establish Rsc1 as a sequence-specific DNA-binding factor that mediates RSC stimulation by polyA motifs at yeast promoters. Overall, this study expands our understanding of the functional diversity of bromodomains and provides insights into the mechanisms governing RSC targeting to chromatin.

## Introduction

The dynamic structure and organisation of eukaryotic chromatin play a major role in regulating nuclear processes such as gene transcription, DNA replication and repair. The fundamental unit of chromatin, the nucleosome, consists of ∼147 base pairs (bp) of DNA wrapped around an octamer of core histones (two copies each of H2A, H2B, H3 and H4). The precise positioning and composition of nucleosomes along the genome are key features of specific gene expression programs (1). The promoters of transcriptionally active genes are characterised by a nucleosome-depleted region (NDR) accessible to transcription factors and to the transcriptional machinery. The NDR is flanked by two nucleosomes referred to as -1 (upstream) and +1 (downstream). The +1 nucleosome is well-positioned and typically located just downstream of the transcription start site (TSS). Nucleosomes are then regularly spaced along the gene body, gradually dissipating with increasing distance from the TSS (2).

Nucleosome positioning is governed by ATP-dependent chromatin remodelers that harness the energy of ATP hydrolysis to translocate DNA along the histone surface (3). In *Saccharomyces cerevisiae*, the only remodeler essential for viability is the RSC complex (4). RSC belongs to the SWI/SWF family of remodeling complexes that generate and maintain NDRs by pushing the +1 and −1 nucleosomes apart (5). Consistent with its role at NDRs (6), RSC preferentially acts on H2A.Z-containing nucleosomes (7), which are enriched at the +1 and -1 positions genome-wide. Cryo-electron microscopy (cryo-EM) structures of yeast (8–12) and mammalian (13–16) SWI/SNF complexes have elucidated their architecture and interactions with the nucleosome. *S. cerevisiae* RSC grips the nucleosome on both sides: the ATPase subunit Sth1 tightly binds nucleosomal DNA and may contact the H4 tail, while the Sfh1 subunit interacts with the acidic patch of the histone octamer. In addition, a DNA-interaction module formed by subunits Rsc1 (or its paralogue Rsc2), Rsc3 and Rsc30 is thought to engage with DNA exiting the nucleosome (10). However, more than 60% of RSC residues, including the DNA interaction module, remain unresolved in cryo-EM structures, leaving key aspects of RSC recruitment poorly understood (17).

RSC recruitment and activity are influenced by chromatin post-translational modifications, particularly histone acetylation. *In vitro*, H3 and H4 acetylation promotes RSC recruitment, while its effect on remodeling activity remains debated (18–20). Bromodomains (BDs), ∼110-amino acid modules that recognise acetylated lysine residues in histone tails and other proteins (21,22), are abundant in RSC and its mammalian ortholog PBAF. RSC contains eight BDs distributed across its Rsc1/Rsc2, Rsc4, Sth1 and Rsc58 subunits. Among these, the BD in Sth1, and to a lesser extent the second BD of Rsc4 (Rsc4-BD2), recognise H3K14 acetylation, which is enriched at TSSs and DNA damage sites (23–26). Rsc4-BD1 binds an acetylated lysine on Rsc4 itself, antagonising Rsc4 binding to H3K14 acetylation (25). In contrast, the acetyllysine-binding activities of Rsc1/Rsc2 and Rsc58 BDs have yet to be characterised.

Evidence suggests that DNA sequence elements also contribute to RSC targeting to chromatin, particularly GC-rich and polyA motifs. A GC-rich motif is recognised by *S. cerevisiae* Rsc3 and Rsc30 *in vitro* and enriched at NDRs sensitive to Rsc3 mutation *in vivo* (27). PolyA motifs stimulate RSC remodeling activity *in vitro* (28), enhance NDR formation in an *in vitro* reconstituted system, and influence the directionality of nucleosome displacement by RSC (29). Genome-wide mapping revealed that both polyA and GC-rich motifs are enriched at NDRs affected by RSC depletion (30). Moreover, the proximity and orientation of paired polyA and GC-rich motifs affected RSC binding and nucleosome displacement (31). While GC-rich motif binding is attributed to Rsc3/Rsc30, the molecular players mediating RSC stimulation by polyA motifs are unknown.

While most RSC studies had been conducted in the model species *S. cerevisiae*, we chose to investigate it in the pathogenic yeast *Candida albicans*. This opportunistic species can cause invasive candidiasis in immunosuppressed and fragile patients, with an estimated worldwide annual incidence of 700000 cases and a mortality rate reaching 40% (32). RSC contains numerous BDs which are seen as promising candidates for the development of small molecule therapeutics that inhibit their acetyl binding pocket, in both human (21,22) and fungi (33–35). This project was based on the premise that RSC is essential for yeast viability and contains multiple BDs. Therefore, we hypothesised that RSC could represent a potential therapeutic target for combating fungal infections, warranting further investigation. We focused our study on Rsc1, a fungi-specific subunit of RSC, bearing 2 BDs, a AT-hook and a bromo-adjacent homology (BAH) domain (**Figure 1A**). *S. cerevisiae* has two paralogues, Rsc1 and Rsc2, mutually exclusive in the complex. The expression of at least one paralogue is necessary for viability, and the presence of BD2 is required (36). In *C. albicans*, where there is no paralogue protein, Rsc1 was suggested to be essential for viability (37). In this context, we investigated Rsc1-BD2 functions, primarily in *C. albicans* and using *S. cerevisiae* as a secondary model.

**Figure 1.**
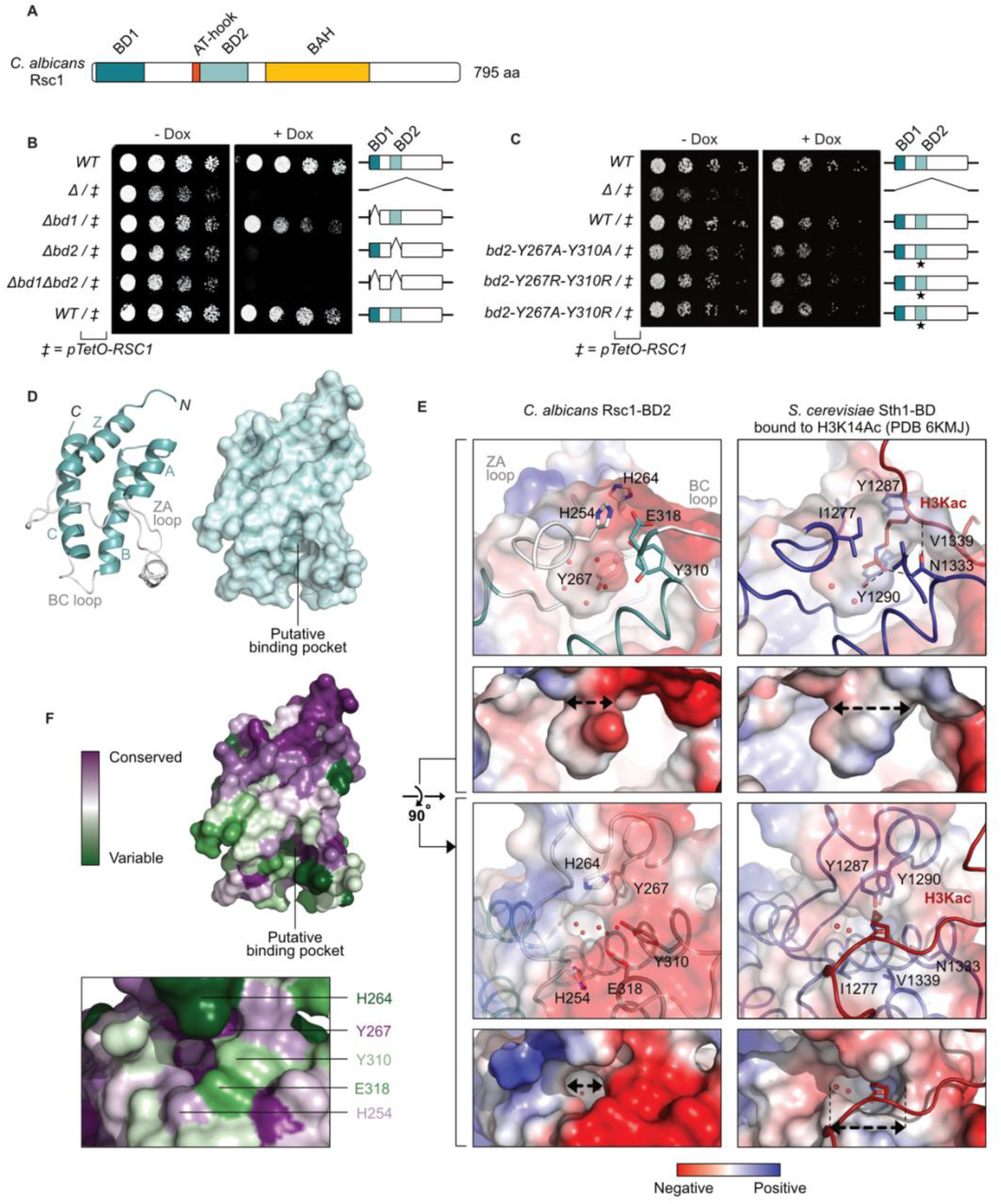
Rsc1-BD2 is essential for *C. albicans* viability but lacks a functional acetyllysine binding pocket. **A.** Domain organisation of *C. albicans* Rsc1 protein. **B-C.** Colony formation assays assessing the viability of strains with conditional *RSC1* gene alterations. One (WT) *RSC1* allele is under the control of a *pTetO* promoter (*pTetO-RSC1*), which is repressed in the presence of doxycycline (Dox), while the other allele is either WT or mutated. Serial dilutions of actively growing cells were plated on media ± Dox and grown for 24 h. Assays were performed in biological duplicates with similar results. **B**. Viability upon deletion of Rsc1 BDs. **C**. Viability upon point mutations of Rsc1-BD2. **D.** Crystal structure of *C. albicans* Rsc1-BD2 at 1.5 Å resolution, revealing the canonical BD fold consisting of four alpha helices (Z, A, B, C), with two of the connecting loops (ZA and BC) defining the putative ligand binding pocket. The four Rsc1-BD2 molecules present in the asymmetric unit were highly similar (aligned structures had a root-mean-square deviation below 0.3 Å for at least 93 Cα). Left: cartoon representation with loops coloured in white. Right: surface representation. **E.** Comparison of the putative binding pocket of *C. albicans* Rsc1-BD2 with the acetyllysine-binding site of *S. cerevisiae* Sth1-BD. *Left:* Close-up view of the Rsc1-BD2 pocket. *Right:* Equivalent views of Sth1-BD bound to an acetylated H3 peptide (PDB 6KMJ, (23)). Cartoon representations are coloured as for panel D for Rsc1; Sth1-BD is in dark blue and the H3 peptide is in red. Water molecules are shown as red spheres. Surfaces are coloured according to their electrostatic potential. Upper panels show the pocket viewed from inside the protein. Lower panels show the pocket viewed from above. **F.** Structure of Rsc1-BD2 coloured according to sequence conservation among fungi. The corresponding sequence alignment is in **Figure S3**.

*Candida albicans* is an opportunistic fungal pathogen that causes invasive infections in immunocompromised patients, with an estimated global incidence of 700,000 cases annually and a mortality rate reaching 40% (32). The need for novel antifungal strategies, together with our previous work on fungal BDs (33–35), led us to explore the BDs of *C. albicans* RSC as potential therapeutic targets. These domains are particularly appealing because RSC is essential for fungal viability and the acetyllysine binding pockets of BDs are highly amenable to small-molecule inhibition (21,22). We focused on Rsc1, a fungal-specific RSC subunit that contains two BDs, an AT-hook and a bromo-adjacent homology (BAH) domain (**Figure 1A**). In *S. cerevisiae*, Rsc1 and its paralogue Rsc2 are mutually exclusive within the RSC complex, with viability dependent on the expression of at least one paralogue and on the presence of BD2 (36). In contrast, *C. albicans* lacks a paralogue and relies solely on Rsc1, which is essential (37). These observations prompted us to investigate Rsc1-BD2 function and its potential as an antifungal target.

In this work, we found that Rsc1-BD2 lacks a functional acetyllysine binding pocket. Instead, together with the adjacent AT-hook and BAH domains, it forms a conserved basic surface that directly interacts with DNA. Single point mutations disrupting this surface are lethal in both *C. albicans* and in *S. cerevisiae*. We further show that Rsc1 DNA binding surface enhances RSC nucleosome remodeling activity both *in vitro* and genome-wide. A *de novo* motif search identified a polyA motif enriched at NDRs in both species, particularly in highly expressed genes. Rsc1 preferentially binds polyA motifs *in vitro*, and nucleosome positioning at NDRs with high polyA motif density is dependent on Rsc1 DNA binding surface. These findings establish Rsc1 as a sequence-specific DNA-binding factor that mediates RSC stimulation by polyA motifs, functioning via an unconventional BD specialised for DNA interaction rather than acetyllysine recognition.

## Results

### Rsc1-BD2 is essential for *C. albicans* viability but lacks a functional acetyllysine binding pocket

To determine whether Rsc1 BDs contribute to viability in *C. albicans*, a diploid yeast, we used a Tet-Off conditional depletion system to knock-down this essential gene (**Figure S1A**) (34). Both *RSC1* alleles were modified at their endogenous loci: one allele was placed under doxycycline (Dox) repression, while the open reading frame (ORF) of the other was either wild-type (WT), fully deleted (Δ), or partially deleted for BD1 (Δbd1), BD2 (Δbd2) or both BDs (Δbd1Δbd2). Colony formation assays (**Figure 1B**) and growth monitoring in liquid culture (**Figure S1B**) showed that all strains grew similarly under permissive conditions (-Dox), while under repressive conditions (+Dox) *RSC1* deletion nearly completely abolished growth, as expected. Notably, deletion of BD2 alone was sufficient to abolish growth, whereas BD1 deletion had only a modest effect. These findings establish Rsc1-BD2 as essential for *C. albicans* viability.

BDs recognise acetylated lysines through a well-characterised binding pocket, in which two conserved residues, an asparagine (N) and a tyrosine (Y), form direct and water-mediated hydrogen bonds with the acetyllysine acetyl group (21,22). In Rsc1-BD2, the tyrosine is conserved (Y267) but the asparagine is replaced by Y310, suggesting a potentially altered ligand-binding activity. To verify binding pocket functionality, we introduced point mutations at these positions. Replacing Y267 with phenylalanine (Y267F), a substitution known to disrupt ligand binding in other yeast BDs (34,38,39), had no effect on growth, even when combined with Y45F, the equivalent mutation in BD1 (**Figure S1C-D**). To further challenge pocket functionality, we introduced more disruptive substitutions by replacing both Y267 and Y310 by either alanine (A) or arginine (R). These mutations also had no impact on viability (**Figure 1C** and **S1E**), despite similar expression levels across all protein variants (**Figure S1F**). These results strongly suggest that a functional BD2 acetyllysine binding pocket is dispensable for Rsc1’s essential activity.

To gain further insights, we determined the crystal structure of *C. albicans* Rsc1-BD2 (**Supplementary Table 1**). The domain adopts a canonical BD fold, with a putative ligand-binding pocket defined by the ZA and BC loops (**Figure 1D**). However, comparison with the BD structure of *S. cerevisiae* RSC subunit Sth1 revealed a markedly narrower pocket entrance in Rsc1-BD2, imposing severe steric restrictions on ligand access (**Figure 1E**). This constriction is due to the bulky side chains of residues H254, Y310 and E318, whereas the corresponding Sth1 residues (I1277, N1333 and V1339) have shorter side chains that define a more accessible binding site. Structural comparisons with additional canonical BDs, both in apo and ligand-bound states, confirmed that the restricted pocket entrance is a unique feature of Rsc1-BD2 (**Figure S2A-B**). Other properties either varied across BDs, such as the electrostatic surface potentials and the configuration of water molecules in the pocket (**Figure S2A**), or were well conserved, such as the buried hydrophobic residues that line the pocket (**Figure S2B**). Given these findings, we examined the conservation of surface-exposed residues in Rsc1-BD2 across fungi. Notably, all pocket residues, including H254, Y310 and E318, exhibited poor conservation, further supporting a loss of acetyllysine-binding functionality (**Figure 1F** and **S3**).

Thus, although Rsc1-BD2 is essential for *C. albicans* viability, its constricted and poorly conserved binding pocket is inconsistent with a conventional acetyllysine reader function. These findings raise the question of what alternative function underlies the essentiality of this domain.

### Rsc1-BD2 forms an essential DNA-binding module with the AT-hook and BAH domains

To gain insights into the structural context of Rsc1-BD2, we used AlphaFold (40) to obtain an atomic model of the full-length *C. albicans* Rsc1 protein. As expected, the model closely aligns with our BD2 crystal structure (root-mean-square deviation [rmsd] of 0.68 Å for 105 Cα atoms; **Figure S4A**). Remarkably, Rsc1-BD2 is predicted to form part of an integrated module that includes the adjacent AT-hook and BAH domains (**Figure 2A**). AT-hooks are small DNA-binding protein motifs characterised by a glycine-arginine-proline (GRP) core flanked by basic lysine (K) or arginine (R) residues, often accompanied by nearby proline (P) residues (41). BAH domains are commonly found in chromatin-associated proteins and often linked to histone binding, including in *S. cerevisiae* Rsc2 (42). In the Rsc1 AlphaFold model, BD2 and BAH are connected by a short linker, forming a crescent-shaped structure with the AT-hook positioned at the hinge. The predicted aligned error (PAE) across this module was low, indicating a well-defined domain arrangement, whereas BD1 and the C-terminal region were predicted to fold independently and to be flexibly tethered. Per-residue confidence estimates, indicated by the predicted local distance difference test (pLDDT) scores (40), were highest for BD2 and BAH, with lower values for the AT-hook and linker, suggesting potential flexibility in these two regions (**Figure S4B**).

**Figure 2.**
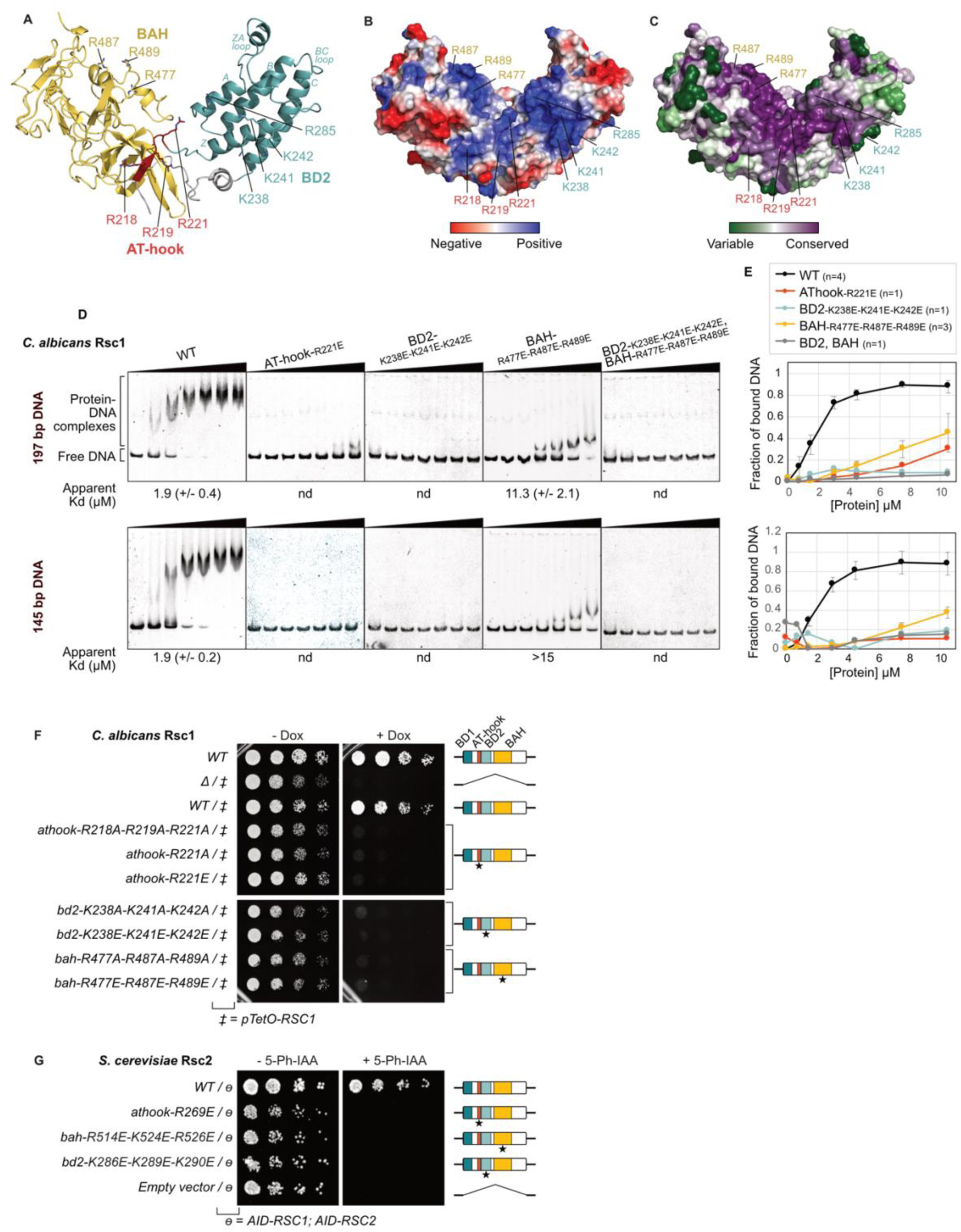
Rsc1-BD2 forms an essential DNA-binding module with the AT-hook and BAH domains. **A-C.** AlphaFold model of *C. albicans* Rsc1 AT-hook, BD2 and BAH domains (residues 213-599). **A.** Cartoon representation. The AT-hook, BD2 and BAH domains are coloured in red, light teal and yellow, respectively. The linker region between BD2 and BAH is coloured in grey. Positively charged surface-exposed residues are shown as sticks. **B.** Surface representation coloured according to the electrostatic potential. **C.** Surface representation coloured according to the sequence conservation among fungi (sequence alignment shown in **Figure S3**). **D.** EMSA of recombinant *C. albicans* Rsc1 proteins (residues 213-599) with 601 DNA probes of 197 and 145 bp on the upper and lower panel, respectively. DNA is at a fixed concentration of 0.15 µM. Recombinant Rsc1 proteins, WT and a series of variants mutated in the basic surface, were used at increasing concentrations of 0, 0.75, 1.5, 3, 4.5, 7.5 and 10.5 µM. Gels were stained to detect DNA. The apparent dissociation constant (*K*_d_) was determined by gel quantification, as shown in panel E (nd: not determined). **E.** Quantifications of EMSAs, including gels shown in panel D and replicates (n). The average fraction of bound DNA is plotted. Error bars indicate the standard deviation. **F**. *C. albicans* colony formation assays using the Tet-Off system for *RSC1* conditional depletion. Serial dilutions of actively growing cells are plated on media ± Dox and grown for 24 h. These assays were performed in biological duplicates with similar results. **G**. *S. cerevisiae* colony formation assays using the AID system for Rsc1 and Rsc2 conditional depletion. Rsc1 and Rsc2 protein were each fused to an AID degron and degraded upon addition of 5-Ph-IAA. A rescue copy of Rsc2, either WT or mutated, is expressed from a pRS316 plasmid. Serial dilutions of actively growing cells are plated on media ± 5-Ph-IAA and grown for 24 h. These assays were performed in biological duplicates with similar results.

Strikingly, the AT-hook-BD2-BAH module displays a large positively charged surface spanning one side of the structure (**Figure 2B**). This basic patch comprises 10 exposed lysine and arginine residues distributed across the three domains (R218, R219, R221 in the AT-hook; K238, K241, K242 and R285 in BD2; and R477, R487 and R489 in BAH). Notably, BD2 residues involved in this basic patch are located on the Z and A helices, far from the putative acetyllysine binding pocket. Conspicuously, this basic surface is strongly conserved across fungi (**Figure 2C** and **S3**), suggesting an important functional role.

We hypothesised that this conserved basic surface mediates DNA binding. To test this, we produced recombinant proteins corresponding to the AT-hook–BD2–BAH module, including the WT version, three variants in which basic residues in either the AT-hook, BD2 or BAH domains were mutated to glutamate (E), and a fourth variant combining BD2 and BAH mutations (**Figure S5A**). Gel filtration chromatography revealed identical elution profiles for all five proteins (**Figure S5B**), while thermal stability measurements using differential scanning fluorimetry (DSF) and NanoDSF yielded comparable melting temperatures (**Figure S5C-D**), confirming that the mutations did not impair protein folding or stability. We then assessed DNA binding using electrophoretic mobility shift assays (EMSAs) with 145- and 197-bp DNA probes containing the 601 positioning sequence (43). The WT module exhibited robust DNA binding, with an apparent dissociation constant (*K*_d_) of 1.9 µM for both DNA probes (**Figure 2D-E**). In contrast, all mutants showed severely impaired DNA binding. Mutations in BD2 (alone or combined with BAH mutations) and in the AT-hook abolished or nearly abolished DNA binding, while mutations in the BAH domain caused a ≥6-fold reduction in binding (*K*_d_ values of 11.3 and >15 µM for the 197 and 145 bp probes, respectively). Thus, the Rsc1 AT-hook–BD2–BAH module directly binds DNA *in vitro,* and this interaction is mediated by the conserved basic surface.

To assess the *in vivo* relevance of this DNA-binding surface, we introduced the same mutations into endogenous *C. albicans RSC1* and examined the effect on viability using our conditional depletion system. In addition to the glutamate substitutions (K/R to E), we also tested milder mutations (K/R to A) and confirmed that all variants were expressed at similar levels (**Figure S6A**). Strikingly, all mutations resulted in severe growth defects (**Figure 2F** and **S6B**), establishing that the basic surface is crucial for Rsc1 function *in vivo*.

Given its strong conservation across fungi (**Figure 2C** and **S3**), we next asked whether this DNA-binding surface plays a similarly important role in *S. cerevisiae*, where the deletion of both *RSC1* and *RSC2* is lethal (36). To address this, we developed a custom Auxin Induced Degron (AID) system, adapted from two recently published AID versions (44,45). Using marker-less CRISPR/Cas9, we added an AID domain to both *RSC1* and *RSC2* in haploid *S. cerevisiae* cells, enabling the simultaneous depletion of both paralogues upon addition of the auxin analog 5-Ph-IAA. We introduced plasmids expressing either WT or mutant versions of Rsc2 and tested their ability to rescue viability following the double depletion (**Figure S6C**). We generated three mutant variants carrying K/R to E mutations in the AT-hook, BD2 or BAH domains, and confirmed that all were expressed at levels comparable to the WT version (**Figure S6D**). Remarkably, none of the mutants rescued growth in the presence of 5-Ph-IAA (**Figure 2G** and **S6E**), indicating that the integrity of the DNA-binding surface is essential for viability.

Together, these findings establish the AT-hook-BD2-BAH module of Rsc1 as a conserved DNA binding factor required for viability in both *C. albicans* and in *S. cerevisiae*.

### Rsc1 basic surface contributes to nucleosome remodeling by the RSC complex

The primary function of the RSC complex is to position nucleosomes at promoter regions. To assess whether the Rsc1 DNA-binding surface contributes to this activity, we first tested whether mutations in this surface affect the composition of the *C. albicans* RSC complex. We immunoprecipitated RSC from strains expressing a Flag-tagged Rsc1 protein, either WT or mutated in BD2 (K238E/K241E/K242E), or from un untagged strain as a negative control. We identified co-purifying proteins by analysing the eluates using mass spectrometry-based quantitative proteomics (**Figure 3A**). Both WT and mutant complexes contained the same 15 proteins with identical estimated stoichiometry (**Figure 3B** and **S7A-B**), consistent with known *C. albicans* RSC subunits. As expected, the complex lacked orthologues of *S. cerevisiae* Rsc3 and Rsc30, which are absent from the *C. albicans* genome, and instead included the *Candida*-specific subunits Nri1 and Nri2 (37). We also detected very low levels of Taf65, the *C. albicans* ortholog of the TFIID subunit Taf8. These findings confirm that mutations in the Rsc1 basic surface do not compromise RSC complex integrity.

**Figure 3.**
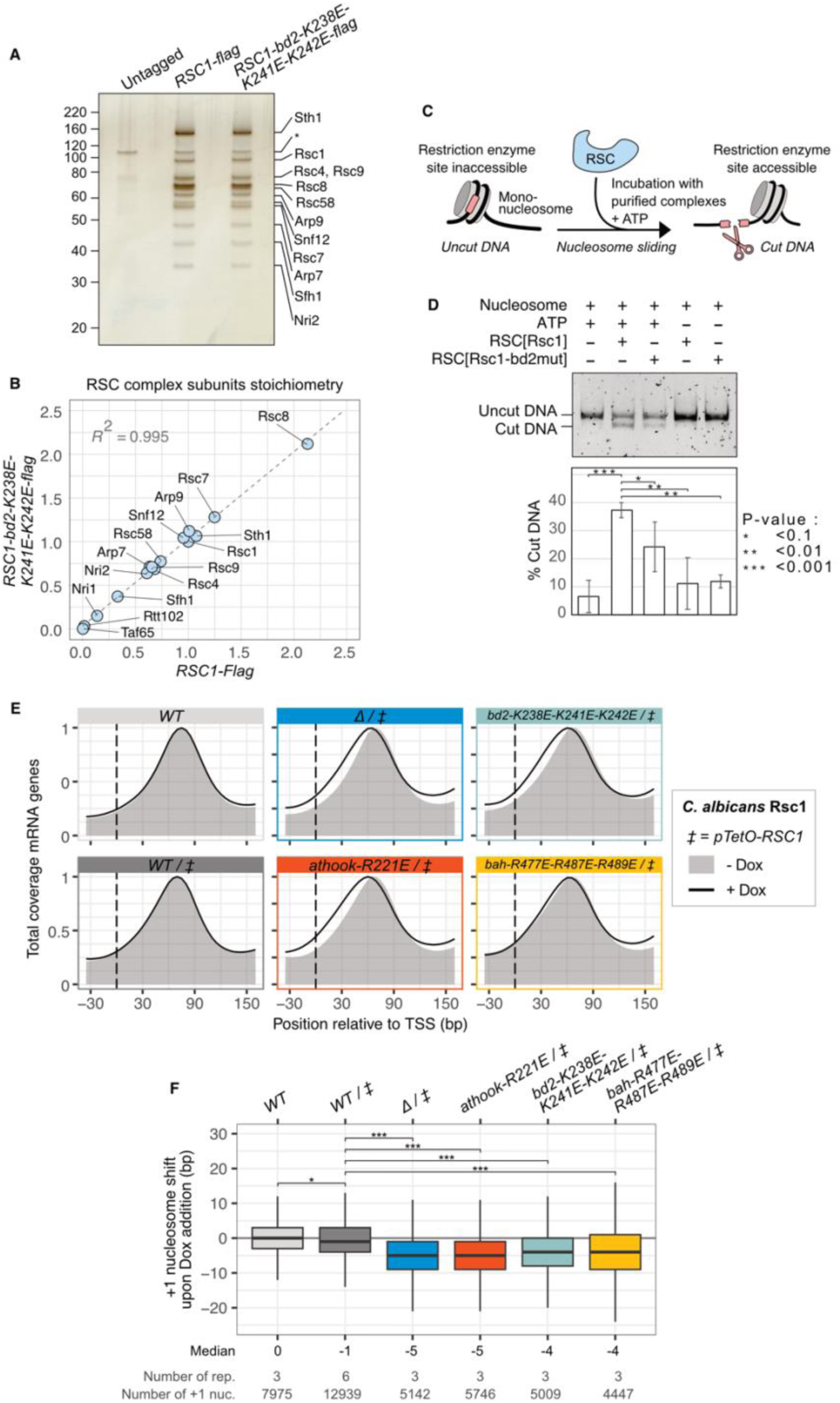
The Rsc1 DNA-binding surface contributes to RSC nucleosome remodeling. **A.** Endogenous RSC complex purified from *C. albicans* by Flag immunoprecipitation. Proteins in immunoprecipitation eluates from *C. albicans* strains, untagged or expressing WT (*RSC1-flag)* or mutant (*RSC1-bd2-K238E-K241E-K242E-flag*) Flag-tagged *RSC1* were detected by silver staining. Protein identities in each band were assigned according to mass spectrometry-based proteomic results and on expected molecular weights. * indicates a non-specific contaminant. **B.** Estimated stoichiometries of RSC subunits, co-purified with Rsc1-flag or Rsc1-bd2-K238E-K241E-K242E-flag, and quantified by mass spectrometry-based proteomics. **C.** Presentation of the Restriction Enzyme Accessibility (REA) assay. An artificial mono-nucleosome shielding a DpnII restriction cut site is used to assess nucleosome remodeling by RSC complexes. In the presence of ATP, the nucleosome is incubated with the complex which, if active, releases the DpnII site for cleavage. DNA then is analysed by gel electrophoresis to visualise cleavage. **D.** REA assay comparing nucleosome remodeling activities of RSC complexes, containing Rsc1-TAP or Rsc1-bd2-K238E-K241E-K242E-TAP. Complexes were purified using TAP purification from *C. albicans* TAP-tagged strains. REA assays were visualised by DNA electrophoresis (top panel: representative gel). The fraction of cut DNA was quantified from triplicate assays (bottom panel: mean +/- standard deviation, P-values from a Dunnett’s multiple comparison test. SDS-PAGE and western-blot analysis of purified complexes are shown in **Figure S7C-D**. **E.** Average position of the +1 nucleosome of protein coding genes in *C. albicans*, determined by MNase-seq. *pTetO-RSC1* is depleted upon addition for Dox, while a second allele, WT or mutated, is constitutively expressed. The coverage over the -30 to +150 bp region relative to the TSS was averaged for the 6221 genes analysed, and normalised between 3 biological replicates. For each strain, the grey shadow and the black line indicate the coverage in permissive (-Dox) or in repressive (+Dox) conditions, respectively. **F.** For each individual MNase-seq sample, the positions the +1 nucleosome was determined for each protein coding gene. The shift of the +1 nucleosome upon Dox addition was calculated for each -Dox and +Dox sample pair. The distribution of this shift is shown as a boxplot, where biological replicates were combined (individual replicates are shown in **Figure S8B**). A negative value indicates that the +1 nucleosome moved closer to the TSS upon Dox addition. The number of observations (number of +1 nuc.) varies between samples as it reflects the number of +1 peaks that could be confidently detected. Each condition was compared to the WT/‡ reference condition using a Dunnett’s test, p-values are indicated as follow, *: <0.01, ***: <0,0001. The medians, number of replicates and the number of +1 nucleosomes included in this analysis are indicated below the plot.

We next assessed whether these mutations affect RSC remodeling activity using an *in vitro* Restriction Enzyme Accessibility (REA) assay. We incubated TAP-purified RSC complexes (**Figure S7C-D**) with recombinant mono-nucleosomes containing a restriction enzyme site that becomes accessible upon RSC activity (**Figure 3C**). As expected, RSC containing WT Rsc1 exhibited ATP-dependent remodeling activity. Notably, this activity was reduced by 35% for RSC containing the Rsc1 BD2 mutant (**Figure 3D**). This suggests that Rsc1-mediated DNA binding enhances RSC remodeling activity, possibly by promoting recruitment to the nucleosome or stabilising RSC-nucleosome interactions.

To investigate whether Rsc1 DNA-binding surface influences nucleosome positioning *in vivo*, we performed genome-wide nucleosome mapping in *C. albicans* using micrococcal nuclease digestion followed by sequencing (MNase-seq). Rsc1 was conditionally deleted or mutated using five of our Tet-Off strains (including WT, deletion, mutations in the AT-hook, BD2 and BAH), a WT strain bearing unmodified *RSC1* alleles was also included (as in **Figure 2F**). MNase-seq was performed in biological triplicates on mock-treated (-Dox) and Dox-treated (+Dox) samples to assess nucleosome positioning in the presence or absence of WT Rsc1. We analysed nucleosomal coverage over the ∼6200 protein coding genes present in the *C. albicans* genome, obtaining high-resolution maps for all samples. Typical profiles were characterised by NDRs flanked by sharply defined -1 and +1 nucleosomes, respectively upstream and downstream of the TSS, followed by phased nucleosomes across the gene bodies (**Figure S8A**). Since yeast RSC primarily functions at NDRs and is critical for +1 nucleosome positioning (27,46), we compared the average nucleosome coverage over the +1 region between mock and Dox-treated samples for each strain (**Figure 3E**). As expected, the +1 nucleosome position remained unchanged in the unmodified WT strain and in the Tet-Off strain expressing WT Rsc1 (*WT/pTetO-RSC1,* +Dox). In contrast, Rsc1 deletion caused a shift of the +1 nucleosome toward the TSS upon Dox treatment (*Δ/pTetO-RSC1*), indicating a reduced accessibility of the NDR. Strikingly, mutations in the AT-hook or BD2 phenocopied this effect, whereas mutations in the BAH domain, despite severely impairing growth, did not significantly alter the +1 nucleosome’s average position.

We then analysed the differential position of +1 nucleosome at each individual gene under mock and Dox treated conditions. For each gene. Negative values indicated that the +1 moved towards the TSS when Dox was added, and the reverse for positive values; zero values indicated no change. For each strain, the distribution of these shift values was presented in **Figure 3F**. This confirmed that the unmodified WT strain was insensitive to Dox, with a distribution centred on zero. The Tet-Off strain expressing WT Rsc1 showed a slightly modified distribution centred on -1 bp, likely reflecting the reduction in Rsc1 protein level upon Dox treatment, which represses one allele out of two. Notably, Rsc1 deletion or mutation in the AT-hook, BD2 or BAH resulted in more pronounced changes, with distributions centred at -5 or - 4 bp. Although this distance was relatively small, it was global for most genes and highly reproducible across replicates (**Figure S8B**). Thus, the depletion of Rsc1 or disruption of its DNA binding surface resulted a widespread repositioning of +1 nucleosomes towards the TSS by 4 to 5 bp, which corresponds to roughly half a helical turn of B-form DNA.

Altogether, our *in vitro* and *in vivo* data demonstrate that the Rsc1 basic surface enhances RSC-dependent nucleosome remodeling. We propose that this DNA-binding interface promotes RSC function by facilitating its recruitment and/or stabilising its association with chromatin.

### An NDR-specific polyA motif is associated with nucleosome architecture and gene expression

The data presented above identified Rsc1 as a DNA binding protein that contributes to RSC activity at NDRs. Given that in *S. cerevisiae*, RSC activity is influenced by DNA sequence both *in vitro* and *in vivo* (27–31), we asked whether specific DNA elements might guide RSC to NDRs via Rsc1-mediated recruitment in *C. albicans*. We performed an unbiased *de novo* motif search in the -300 to +100 bp region around the TSS (corresponding to the NDR) of all *C. albicans* protein coding genes. This analysis revealed a 15 bp A-rich motif composed of an initial G or A followed by a stretch of A bases with high information content (**Figure 4A**). This motif was specifically enriched in the -300 to +100 bp interval compared to adjacent regions (**Figure S9A**).

**Figure 4.**
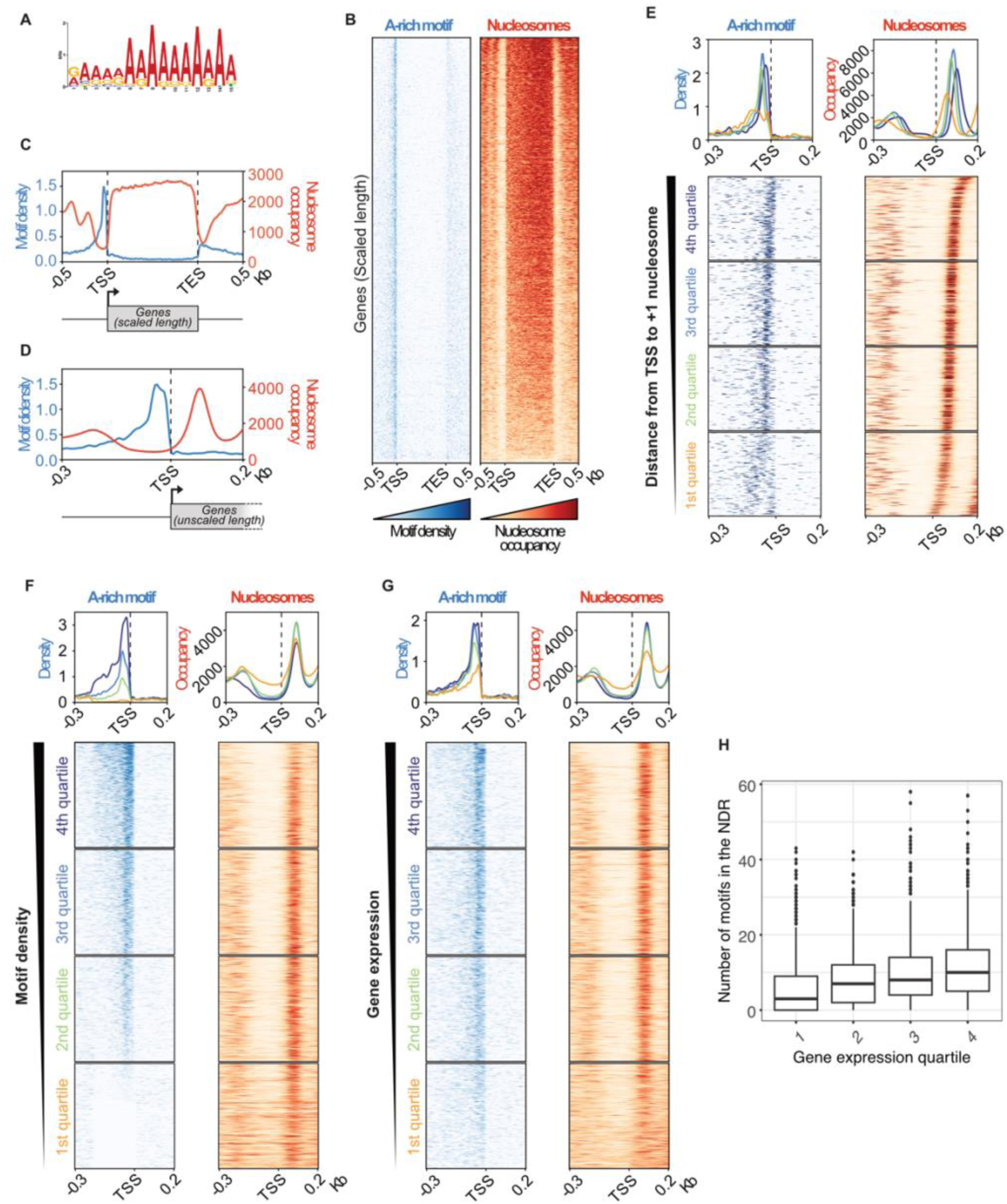
An NDR-specific A-rich motif is associated with nucleosome architecture and gene expression in *C. albicans*. **A.** A-rich Motif identified *de novo* in the NDR region (-300 to +100 bp around the TSS) of *C. albicans* protein coding genes. **B.** Heatmaps of the A-rich motif density (left side, coloured blue) and nucleosomal occupancy (right side, coloured orange) over all protein coding genes (6221 genes). Gene length is scaled from TSS to TES, with 0.5 Kb flanks. The nucleosomal occupancy is from our MNase-seq data (WT sample without Dox). **C.** Average profiles of motif density (blue curve, number of motifs) and nucleosomal occupancy (orange curve, RPKM) over all protein coding genes, corresponding to heatmaps shown in panel **A**. **D.** As in panel **C** but gene length is unscaled, genes are aligned to TSS with -0.3 and 0.2 Kb flanks. **E.** Motif density and nucleosomal occupancy over genes sorted and stratified by the distance from the TSS to the +1 nucleosome (3785 genes for which the +1 has been confidently mapped). Profiles (top) and heatmaps (bottom) are shown for each quartile. **F.** As in panel **E** but for all protein coding genes, sorted by the number of motifs present upstream of the TSS (from -0.2 Kb to TSS), and stratified by quartiles. **G.** As in panel **E** but genes are sorted and stratified by gene expression, based on RNA-seq data (47,48). **H.** Boxplot of the number of motifs present upstream of the TSS (from - 0.2 Kb to TSS) of all protein coding genes stratified by gene expression.

We analysed the distribution of this motif along the genes and flanking regions, and compared it with nucleosome occupancy determined from our MNase-seq data (**Figure 4B-D**). Strikingly, the motif is present in the vast majority of gene promoters (**Figure 4B**). It is highly focused at the NDRs, where nucleosomal occupancy is the lowest, with modest enrichment after the Transcription End Site (TES) and a clear exclusion from gene bodies (**Figure 4B-C**). Closer inspection revealed that the motif peaks between the -1 nucleosome and the TSS, with a sharp drop in density precisely at the TSS (**Figure 4D**). Thus, we identified a polyA motif specifically enriched at NDRs and widespread across protein coding genes in *C. albicans*.

To refine these observations, we analysed the distribution of the motif relative to the +1 nucleosome. For this, we selected genes with well-defined +1 nucleosome positions, ranked them by TSS:+1 distance, stratified them into quartiles, and plotted motif and nucleosome distributions (**Figure 4E**). Notably, the motif position mirrors that of the +1 nucleosome, suggesting a functional link. In the first quartile, where the +1 is closest to the TSS, we observed two motif populations, one just upstream of the TSS (-15 bp) and another more diffusely localised further upstream (-80 bp). We then ranked all protein coding genes by the number of motifs present in the 200 bp region upstream of the TSS. This revealed that a higher motif density correlates with lower nucleosome occupancy and a wider NDR, due to a more distal -1 nucleosome (**Figure 4F**).

To evaluate whether motif density correlates with gene expression levels, we used publicly available RNA-seq data (47,48) and ranked genes by transcript abundance (quantified in Fragments Per Kilobase Million or FPKM). Highly expressed genes exhibited higher motif density, more precisely positioned +1 nucleosomes, lower nucleosome occupancy within the NDR, and wider NDRs (**Figure 4G**). This positive correlation between gene expression and motif density was further supported by the distribution of motif counts across gene expression quartiles (**Figure 4H**).

Finally, we asked whether these observations extend to *S. cerevisiae*. Applying our motif search strategy to *S. cerevisiae* NDRs revealed a polyA motif very similar to that found in *C. albicans* (**Figure S9B**), which was also specifically enriched in NDRs and regions of low nucleosome occupancy (**Figure S9C-D**). Furthermore, as in *C. albicans*, a higher motif density correlates with increased gene expression and wider NDRs (**Figure S9E**).

Thus, *C. albicans* promoters are enriched for polyA motifs, whose distribution and density are strongly associated with NDR architecture and gene expression levels. This pattern is conserved in *S. cerevisiae*, suggesting a fundamental role for these motifs in chromatin organisation and transcriptional regulation.

### Rsc1 action is driven by A-rich sequences

As shown above, conditional deletion or mutation of Rsc1 caused the +1 nucleosome to shift toward the TSS upon Rsc1 conditional deletion and mutation (**Figure 3E-F**). We next sought to determine whether the presence of the polyA motif influences Rsc1 activity by analysing the shift of the +1 nucleosome as a function of motif density. To achieve this, we categorised genes into deciles based on the NDR motif density. We then compared the average +1 nucleosome position between mock and Dox-treated samples in the top and bottom deciles (**Figure 5A**). Genes in the top decile displayed a clear shift of the +1 nucleosome upon Rsc1 deletion and mutation of the DNA-binding surface, whereas those in the bottom decile showed little or no shift. Since the motif density and gene expression are related (**Figure 4G-H**), we expected that the impact of Rsc1 mutations would also depend on gene expression. Indeed, when genes were grouped into expression-based quartiles (**Figure 5B**), highly expressed genes showed a clear +1 shift upon Rsc1 alteration, whereas poorly expressed genes showed a poorly defined +1 peak and were largely unaffected. These findings show that the effect of Rsc1 on the +1 nucleosome is enhanced by the presence of A-rich motifs in the NDR.

**Figure 5.**
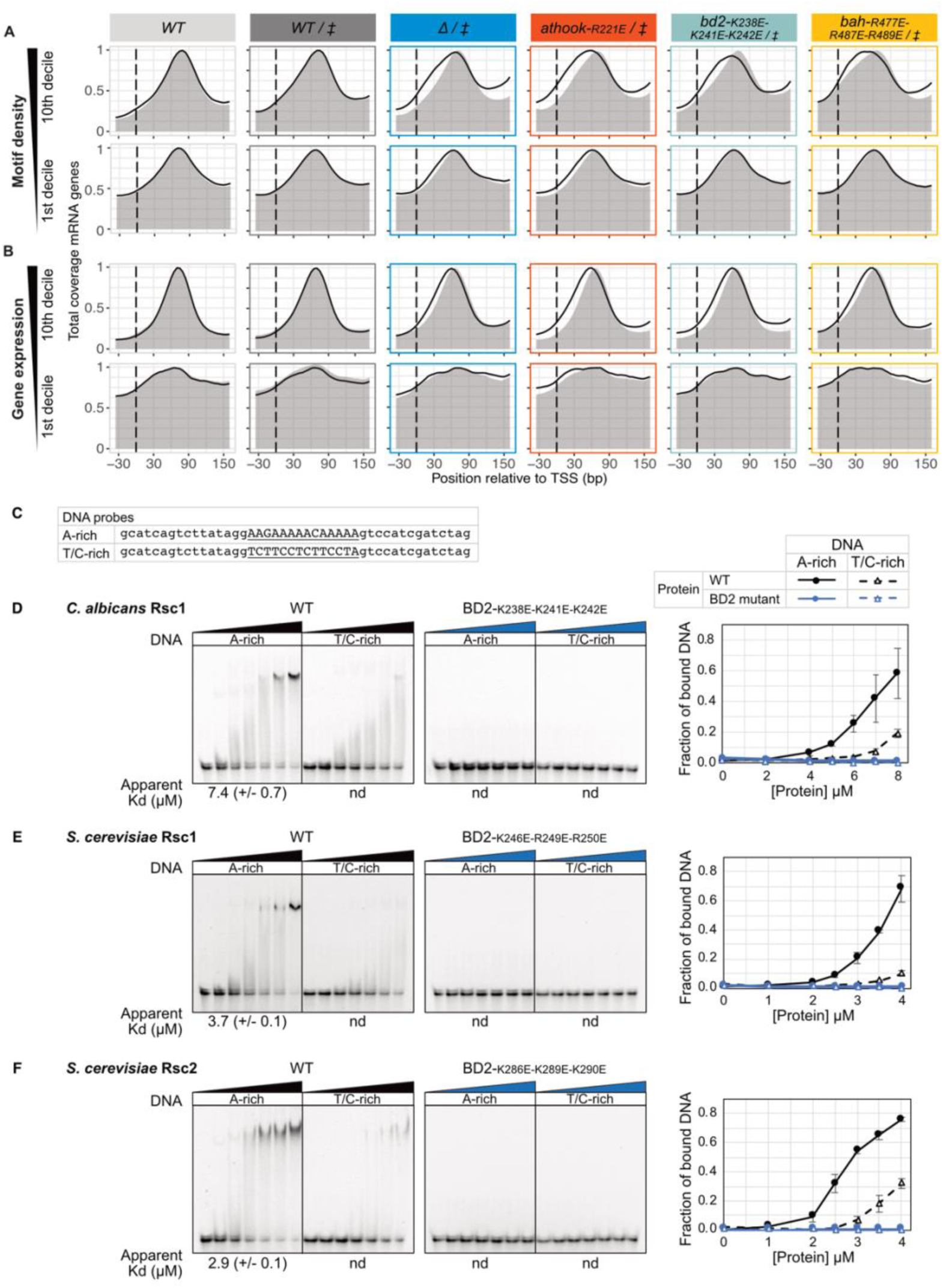
Rsc1 action is driven by A-rich sequences. **A.** *C. albicans* protein coding genes are stratified into deciles, based on the number of motifs present upstream of the TSS. For the 10^th^ and 1^st^ deciles (602 and 603 genes with high and low motif density, respectively), the average position of the +1 nucleosome is shown. The MNase-seq coverage over the -30 to +150 bp region relative to the TSS was averaged for the genes analysed, and normalised between replicates. For each strain, the grey shadow and the black line indicate the coverage in permissive (-Dox) or in repressive (+Dox) conditions, respectively. **B.** As for panel **A**, but genes are stratified into deciles, based on gene expression (47,48). **C.** DNA probes used for EMSAs in panels **D-F**. The A-rich probe matches the motif found enriched in the NDR region (underlined sequence), while the T/C-rich probe contains an unrelated T/C-rich sequence, flanked by fixed sequences. These two of 44 bp probes were 5’ Cy5-labelled and used at a fixed concentration of 20 nM, in the presence of unlabelled non-specific DNA competitor. **D.** EMSA performed with recombinant *C. albicans* Rsc1 (Calb_Rsc1 213-599) WT or mutated in BD2 basic surface (K238E-K241E-K242E) used at increasing concentrations of 0, 2, 4, 5, 6, 7, and 8 µM. Gels and quantifications are shown on the left and right panels, respectively. For quantifications (n=2 for the WT, n=1 for the mutant), the average fraction of bound DNA is plotted, error bars indicate the standard deviation. The Kd was determined from quantifications (nd: not determined). **E.** As for panel **D**, but using recombinant *S. cerevisiae* Rsc1 (Scer_Rsc1 221-596) WT or mutated in BD2 (K246E-R249E-R250E), at 0, 1, 2, 2.5, 3, 3.5 and 4 µM. **F.** As for panel **D**, but using recombinant *S. Cerevisiae* Rsc2 (Scer_Rsc2 261-641) WT or mutated in BD2 (K286E-K289E-K290E), at 0, 1, 2, 2.5, 3, 3.5 and 4 µM.

To test whether Rsc1 preferentially binds DNA containing this polyA motif *in vitro*, we performed EMSAs using two different 44 bp probes, containing either a polyA or a T/C-rich sequence, both flanked by identical random sequences (**Figure 5C**). *C. albicans* Rsc1 bound the polyA probe, forming a well-defined protein-DNA complex, with an apparent *K*_d_ of 7.4 µM (**Figure 5D**). In contrast, binding to the T/C-rich probe was significantly weaker, with no measurable *K*_d_. As a control, we used an Rsc1 variant mutated in BD2 (BD2_K238E-K241E-K242E_), which failed to bind either probe, confirming that DNA binding occurs via Rsc1’s conserved basic surface. The apparent *K*_d_ measured for the polyA probe (**Figure 5D**) was higher than that obtained with the 601 probes (**Figure 2D**). This difference likely reflects variations in probe length and competitor DNA concentration, as the 145- and 197-bp 601 sequences were tested without a competitor, whereas the shorter fluorescently labelled polyA probe was analysed in the presence of an unlabelled, non-specific DNA competitor.

Since the Rsc1 basic surface is conserved and functional in *S. cerevisiae* (**Figure 2C**, **G**), we next evaluated the DNA binding properties of *S. cerevisiae* Rsc1 and its paralogue Rsc2. We purified recombinant WT and mutant forms of both proteins (**Figure S5A**). EMSAs revealed that both Rsc1 and Rsc2 bound to DNA with a strong preference for the polyA probe, exhibiting apparent *K*_d_ values of 3.7 µM and 2.9 µM, respectively, while showing only weak binding to the T/C-rich probe (**Figure 5E-F**). Mutation of the basic surface abolished DNA binding in both proteins, demonstrating that, as in *C. albicans* Rsc1, DNA binding is mediated by this conserved surface.

To further characterise the DNA binding specificity of *C. albicans* Rsc1 and *S. cerevisiae* Rsc2, we used DNA affinity purification sequencing (ampDAP-seq) (49) as an orthogonal approach. This technique enables the determination of intrinsic DNA-binding specificity under *in vitro* conditions, eliminating potential biases from protein partners or chromatin context. Proteins were produced using a cell-free protein expression system and incubated with an exogenous DNA library generated from genomic DNA of *Arabidopsis thaliana*, an unrelated species. Interestingly, ampDAP-seq analysis identified a polyA motif for *S. cerevisiae* Rsc2 (**Figure S9B**), which closely resembled the NDR-specific motif detected in *C. albicans* and *S. cerevisiae* (**Figure 4A**). Technical limitations prevented us from detecting significant motif enrichment for *C. albicans* Rsc1, possibly due to its slightly lower binding affinity for the polyA probe compared to *S. cerevisiae* Rsc2 (7.4 and 2.9 µM, respectively, **Figure 5D**, **F**).

Taken together, these findings establish *C. albicans* Rsc1 and *S. cerevisiae* Rsc1 and Rsc2 as DNA-binding proteins with a strong preference for polyA sequences. Furthermore, we showed that in *C. albicans*, the presence of polyA motifs in the NDR influences nucleosome remodeling by RSC in a Rsc1-dependent manner.

### Rsc1-BD2 underwent a unique functional switch during fungal BD evolution

Our results suggest that Rsc1-BD2 underwent a significant functional transition during molecular evolution, from canonical acetyllysine recognition to DNA binding. To place this shift in a broader context, we analysed all BD sequences from 15 phylogenetically diverse yeast species (50). We focused on the conservation of residues critical for acetyllysine recognition and DNA binding (**Figure 6A**), which we represent as heatmaps (**Figure 6B-C**). The results, which extend our prior analysis of Rsc1-BD2 sequence conservation (**Figure 1F**, **2C** and **S3**) beyond the *S. cerevisiae* and *C. albicans* lineages, reveal that the DNA-binding basic residues in Rsc1-BD2 are conserved in all species examined (**Figure 6B**). This suggests that Rsc1 DNA-binding activity arose before the divergence of the *Ascomycota* phylum, which encompasses all these species. Of the 15 Rsc1-BD2 sequences analysed, 9 lack one or both of the critical acetyllysine binding residues, while 6 retain both (**Figure 6C**). The latter are distributed in a patchwork-like manner across various lineages, suggesting either functional inactivation of the binding pocket by additional mutations or fluctuating selective pressure on acetyllysine recognition over evolutionary time. Further studies are needed to distinguish between these possibilities.

**Figure 6.**
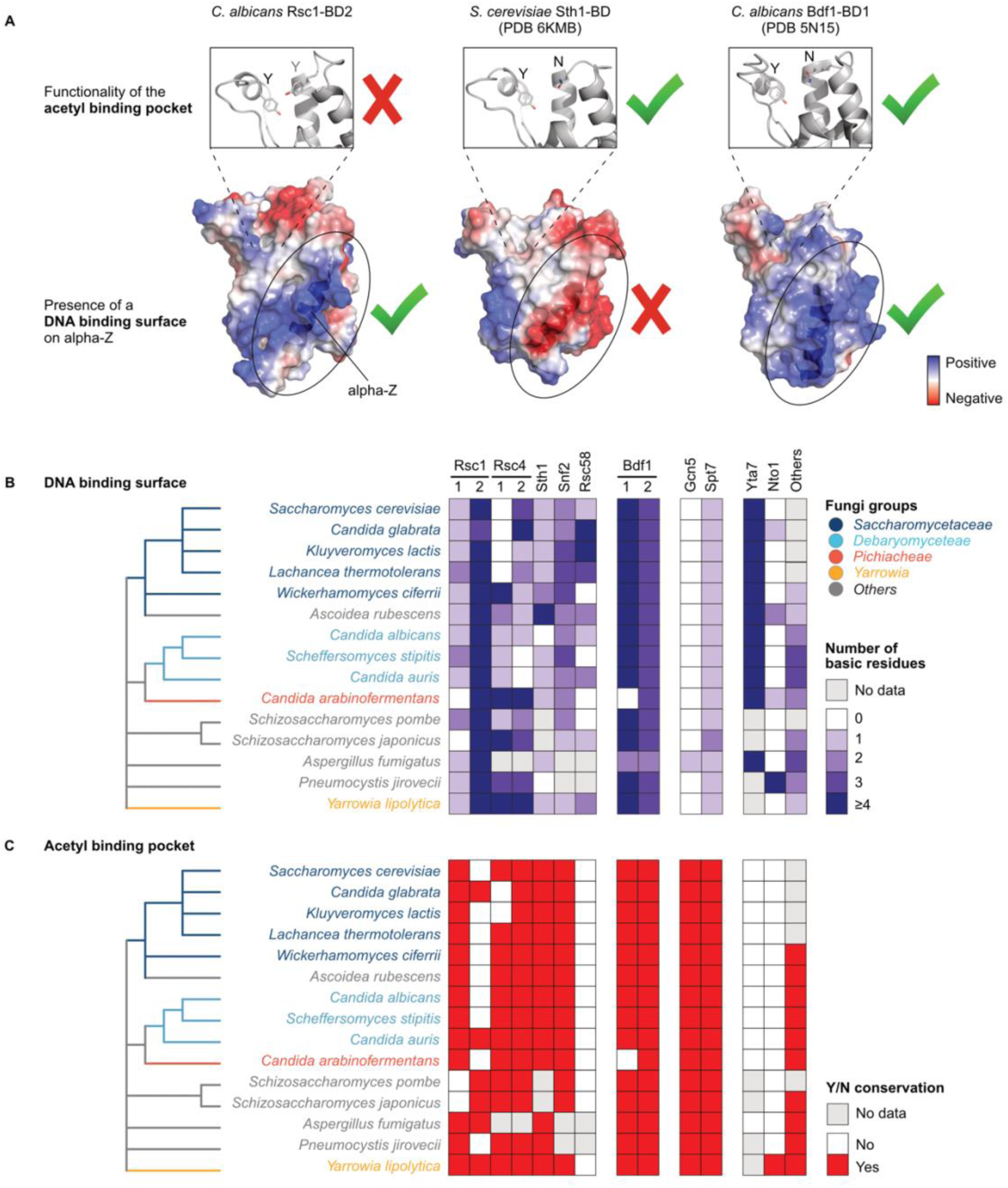
Phylogenetic diversity of BD structural determinants in fungi. **A.** *C. albicans* Rsc1-BD2, *S. cerevisiae* Sth1-BD and *C. albicans* Bdf1-BD1 crystal structures highlighting the key features for the functionality of the DNA-binding surface and acetylated lysine-binding pocket. **B-C.** Heatmaps showing the conservation of key residues for the DNA-binding surface function (**B**) and acetylated lysine-binding pocket (**C**). 15 yeast species were selected as representative of fungi. Their BD sequences were analysed to determine the conservation of key residues for DNA-binding surface and acetylated lysine-binding pocket function. Protein ID from which BD sequences were extracted are presented in **Table S5**. Phylogenetic trees represent the relationship between species and have been extracted from NCBI Taxonomy (50).

Across the broader family of yeast BDs, most domains retain the two key acetyllysine binding residues, with notable exceptions including Yta7 Rsc58 and Nto1 (**Figure 6C**). Yta7-BD was recently shown to interact with the nucleosome in a unique manner, contributing to H3 binding independently of H3 acetylation (51). Interestingly, Yta7-BD features a strongly basic surface resembling that of Rsc1-BD2 (**Figure 6B**), raising the possibility that direct DNA binding may contribute to the Yta7-nucleosome interaction. Rsc58 is an essential RSC subunit in *S. cerevisiae*. Although resolved in RSC cryo-EM structures, the precise function of Rsc58-BD remains unclear. The lack of key acetyllysine binding residues (**Figure 6C**) points to a non-canonical role. Finally, in almost all species analysed, Bdf1-BD1, and to a lesser extent Bdf1-BD2, exhibit numerous basic residues that align with those in Rsc1-BD2 (**Figure 6A**, **B**), while preserving canonical acetyllysine binding features (**Figure 6A**, **C**). Given that the human homologs BRDT and BRD4 bind DNA *in vitro* (52,53), fungal Bdf1-BDs may also possess dual DNA and acetyllysine binding capability. This analysis highlights DNA binding surfaces that aligned to that of Rsc1-BD2, it does not exclude other basic surfaces that may locate elsewhere on the fungal BDs.

These findings reveal a remarkable functional divergence within the BD family. While most BDs maintain their acetyllysine reader function, Rsc1-BD2 appears to have lost this capacity and instead evolved a specialised DNA-binding role that is deeply conserved in *Ascomycota*. Rsc1-BD2 thus emerges as a unique BD whose repurposing from acetyllysine recognition to DNA interaction offers insights into the evolutionary plasticity of BDs and the functional specialisation of protein domains in fungi.

## Discussion

### Rsc1-BD2 unveils a novel type of non-canonical BD

Our study identifies Rsc1-BD2 as a noncanonical BD with a function distinct from acetyllysine recognition. Structural and evolutionary analyses revealed that the predicted acetyllysine binding pocket is largely dispensable for *C. albicans* viability and is characterised by a constricted entrance that is incompatible with acetyl recognition, defined by residues of low evolutionary conservation (**Figure 1**). Instead, we demonstrate that Rsc1-BD2, together with the adjacent AT-hook and BAH domains, constitutes a conserved basic module that directly engages DNA. This DNA-binding activity is essential for viability in both *C. albicans* and *S. cerevisiae* (**Figure 2**) and is critical for the nucleosome remodelling function of RSC (**Figure 3**). A comprehensive phylogenetic analysis of fungal Rsc1-BD2 homologues suggests that the potential for acetyl-lysine binding has fluctuated throughout evolution, while the DNA-binding interface remains highly conserved, underscoring a specialised role for Rsc1 -BD2 in DNA binding (**Figure 6**).

BET proteins are well-characterised acetylated nucleosome readers. Yet human BET BDs also interact with nucleic acids (52–57). A pioneering study revealed that BRDT-BD1 binds DNA via a basic alpha-Z helix (52), corresponding to the DNA-binding patch we identified in Rsc1-BD2. This interface is distinct from the histone-binding pocket, enabling dual recognition of DNA and acetylated nucleosomes (52). Similarly, BRD4 tandem BDs associate with RNA (54) and DNA (55), enhancing chromatin occupancy and cofactor activity, while BRD3 BDs interact with non-coding RNA (56). Given the conserved alpha-Z helix, we propose a similar DNA-binding interface in fungal BET Bdf1 (**Figure 6**), though its nucleic acid-binding capacity and functional relevance remain untested. Whether Rsc1 also interacts with RNA warrants further investigation.

Nucleic acid binding has also been reported for BDs in human SWI/SNF complexes. The ATPase subunits BRG1/SMARCA4 and BRM/SMARCA2 harbour a conserved C-terminal AT-hook adjacent to a BD, forming a DNA-binding module with moderate AT-rich sequence preference (58–60). BRM-BD binds both DNA and H3K14-acetylated histones, suggesting a competitive binding mode (58). Although the AT-hook–BD arrangement in BRM resembles that of Rsc1, their DNA-binding interfaces are distinct. Rsc1-BD2 engages DNA via a basic patch on the alpha-Z helix, spatially separated from the non-functional acetyllysine pocket (**Figure 2A**). In contrast, BRM-BD interacts with DNA via the ZA-loop, overlapping with the histone-binding site, supporting a model in which DNA competes with H3K14-acetyl recognition (58). Consistently, we did not identify significant basic surfaces matching that of Rsc1-BD2 in the fungal SWI/SNF ATPases, Sth1 and Snf2 (**Figure 6**). The PBRM1 subunit of the human PBAF complex, which contains six BDs and two BAH domains, also engages nucleic acids. BD2 and BD4 associate with double-stranded RNA (61), a mechanism proposed to enhance H3K14-acetyl recognition and chromatin association. Notably, PBRM1-BD2 and BD4 interact with RNA via distinct surfaces (61), neither of which resembles the DNA-binding interface of Rsc1-BD2.

Collectively, multiple human BDs interact with DNA or RNA through distinct basic surfaces, often modulating acetylated histone binding and chromatin targeting. Rsc1-BD2 emerges as unique, functioning primarily, if not exclusively, through its DNA-binding interface, while its acetyllysine pocket remains inactive. Moreover, it operates in concert with an adjacent AT-hook and BAH domain, a structural arrangement not observed in other BDs.

### Rsc1 drives RSC activity at A-rich promoters

In the *C. albicans* genome, as well as in *S. cerevisiae*, we identified the presence of a polyA motif using an unbiased *de novo* motif search. This motif is specific to NDRs and is present in the vast majority of genes (**Figure 4**). More specifically, the polyA motif we identified consists of a G followed by a stretch of A nucleotides. We observed that the distribution and density of this motif were closely related to NDR architecture and gene expression.

The existence of this motif had already been reported in *S. cerevisiae* promoters, where it was referred to as the GA element. It was first found to correlate with NDR position, suggesting an interplay with chromatin remodelers (62). Another study showed that this motif was enriched in *S. cerevisiae* promoters lacking a TATA box (the majority of promoters), and suggested that it plays a direct role in transcription, independently of nucleosome positioning (63). PolyA motifs were later shown to facilitate the activity of the RSC complex, both *in vitro* (28,29) and *in vivo* (30,31), supporting a direct interplay between polyA and RSC.

Here, we demonstrate that *C. albicans* Rsc1 and *S. cerevisiae* Rsc1/Rsc2 directly and specifically bind to the polyA motif *in vitro*, and we further show that Rsc1-dependent nucleosome remodeling is conditioned by the presence of this motif in *C. albicans* (**Figure 5**). We therefore propose that the polyA motif present at yeast NDRs is recognised by the Rsc1 protein, through its DNA-binding module, which mediates the recruitment of the RSC complex to promoter regions and contributes to NDR maintenance. This model is compatible with a potential additional role of the polyA motif on the transcriptional machinery, perhaps mediated by other DNA binding proteins.

In *S. cerevisiae*, a GC-rich motif was also found at promoters, in addition to the polyA motif; it is recognised by the Rsc3/30 subunits and has also been correlated with RSC activity (27,31). In *C. albicans*, since Rsc3/30 are absent, RSC sequence specificity may depend entirely on Rsc1.

Cryo-EM structures of the RSC complex have provided significant insights into its architecture, yet they remain incomplete with respect to key functional domains (8–10). The available models include only a short C-terminal region of the Rsc1 subunit. Based on cryo-EM and crosslinking mass spectrometry data, Rsc1 has been proposed to be part of a DNA-interaction module and to engage with DNA exiting the nucleosome (10). Further structural studies will be key to determine how the DNA binding module of Rsc1 engages with DNA, relative to other RSC core modules and in the context of the nucleosome.

SWI/SNF complexes are highly conserved in eukaryotes, however the Rsc1 protein is specific to the yeast RSC complex. While mammalian SWI/SNF complexes contain multiple DNA-binding domains distributed across various subunits, none display clear sequence specificity. Therefore, whereas the RSC DNA-binding module may serve to anchor the complex at promoters during remodeling cycles in yeast, mammalian SWI/SNF complexes likely rely on alternative mechanisms for chromatin retention, including cooperative interactions with multiple transcription factors, recognition of histone modifications, and engagement with non-coding RNAs (17).

In conclusion, this study enhances our understanding of the functional diversity of BDs and provides new insights into the mechanisms that direct the RSC complex targeting to chromatin.

## Material and methods

All yeast strains used in this study are listed in **Supplementary Table 2**. Plasmids are listed in **Supplementary Table 3**. Oligonucleotides and gene fragments are listed **Supplementary Table 4**.

### Yeast strains and genome editing

*C. albicans* strains were derived from the diploid SN152 strain (64). The endogenous of *RSC1* locus was modified by genomic integration of cassettes flanked by ∼500 bp homologous regions. Transformation cassettes were obtained by linearising plasmids. *C. albicans* was transformed by a lithium acetate procedure (65). Genomic integrations were verified by PCR and by Sanger sequencing of the modified loci.

Conditional depletion of one *RSC1* allele was achieved by integrating a tetracycline-regulatable cassette upstream of *RSC1* open reading frame (ORF). As previously described (34), this cassette contained a selective marker (*ARG4*), a transactivator (*TetR-VP16*) and seven tandem Tet-operator elements. In addition, three repeats of the v5 epitope tag were N-terminally fused to Rsc1 (cassette cloned in the pJG453 plasmid).

Deletion of one *RSC1* allele was achieved by replacing *RSC1* ORF by a *NatMX6* cassette (plasmid pJG478), conferring nourseothricin resistance.

The series of strains bearing mutations or partial deletions in the *RSC1* coding sequence were obtained by reintegration of a cassette at the *rsc1Δ::NatMX6* locus, which encoded the modified Rsc1 proteins C-terminally fused to three repeats of the Flag epitope tag and a selective marker (*LEU2*); corresponding plasmids are listed in **Supplementary Table 2**).

The Rsc1-TAP strain was obtained by integrating a tandem affinity purification tag (TAP-tag) C-terminally fused to Rsc1 and the coding sequence was followed an *ARG4* marker (plasmid pJG502). The TAP-tag comprised a calmodulin-binding peptide (CBP), a Tobacco etch virus (TEV) protease cleavage site and two repeats of protein A. The strain expressing Rsc1-bd2-K238E-K241E-K242E-TAP was obtained by replacing *RSC1* ORF by a mutated *RSC1* ORF fused to a TAP-tag and followed by *ARG4* (plasmid pJG571).

Plasmids containing the *C. albicans* transformation cassettes were assembled using NEBuilder HiFi DNA Assembly Master Mix (NEB E2621L), with a backbone sequence derived from pCR-Blunt II-TOPO (Invitrogen 450245). Plasmids were validated by Sanger sequencing.

*S. cerevisiae* strains were derived from BY4741 (66). Conditional simultaneous depletion of *S. cerevisiae* Rsc1 and Rsc2 was achieved by adapting an auxin-inducible degron (AID) system previously described in fission yeast (45). Both endogenous proteins were N-terminally tagged with three repeats of *Arabidopsis thaliana* IAA17^71-106^ (thereafter termed AID), an XTEN16 linker, three repeats of the v5 or HA epitope tag for Rsc1 and Rsc2, respectively, and a short linker (GGGGS). Sequences encoding these tags were added to *RSC1* and *RSC2* loci by two successive rounds of marker-less CRISPR/Cas9 genome editing, using pWS158 (Addgene 90517) and pWS171 (Addgene 90518) as Cas9 expression vectors, as previously described (67). For each *RSC1* and *RSC2*, a protospacer adjacent motifs (PAM) sequence within 30 nucleotides of the start codon was selected and a guide RNA (gRNA) was designed using the Benchling CRISPR tool (http://www.benchling.com). gRNAs were cloned into the pWS082 (Addgene 90517) using annealed oligonucleotides by Esp3I Golden Gate assembly. Repair templates for homology-directed repair were designed as ∼700 bp fragments to insert the N-terminal tags, using 80-120bp homology flanks. The gRNA sequences were disrupted by insertion of the tag for *RSC1*, and by introducing a non-coding mutation upstream of the start codon for *RSC2*. Repair templates were purchased as double stranded DNA fragments (gBlocks) from Integrated DNA Technologies Inc. and were PCR amplified to transform into yeast. *S. cerevisiae* cells were transformed with EcoRV-digested gRNA vector (200 ng), Esp3I-digested pWS158 or pWS171 (100 ng) and >2µg of repair template. The gRNA-containing and Cas9-containing fragments were designed to gap-repair in yeast to obtain a vector expressing both Cas9 and the gRNA (67). Yeast cells were transformed using a lithium acetate method (68). Strains were validated by PCR and by Sanger sequencing of the modified loci. The pBRT1 plasmid (69) containing OsTIR1, a NatMX6 cassette and *HIS3*-flanking sequences for genomic insertion was modified to introduce the OsTIR1^F74A^ mutation previously described (45). The resulting pJG544 plasmid was linearised using PmeI and transformed, with correct integration confirmed by PCR. pRS316 (70) was used to create a series of plasmids expressing WT or mutated Rsc2. A genomic region of ∼400 bp upstream of the *RSC2* ORF was used as a promoter and three repeats of Flag epitope were N-terminally fused to Rsc2 (**Supplementary Table 3**).

### Yeast growth assays

*C. albicans* growth assays were performed by growing cells overnight in Synthetic Defined (SD) yeast medium, at 30 °C with agitation. Cultures were then diluted in SD or in SD containing 50 µg/mL doxycycline (SD+Dox) and grown for 5 h (reaching an OD_595nm_ of ∼0.4). Cells were washed and diluted to an OD_595nm_ of 0.2 in H_2_O. For colony formation assays, cells were spotted on SD or SD+Dox agar plates in a 4-fold dilution series, starting from an OD_595nm_ of 0.2, and incubated at 30 °C for 1 day before imaging. For growth assays in liquid medium, cells were prepared similarly, cells diluted in H_2_O were further diluted to an OD_595nm_ of 0.1 in SD or SD+Dox, in a 96-well plate; OD_595nm_ was monitored for 24 h at 30 °C, using a Multiskan FC microplate reader (Thermo Scientific).

*S. cerevisiae* growth assays were performed as for *C. albicans*, with the following differences: cells were grown overnight in SD without uracil (SD-ura), diluted and grown in SD-ura from OD_595nm_ of 0.1 to 0.4-0.8, washed and diluted in H_2_O before being grown on agar plate or in liquid medium containing either SD-ura of SD-ura supplemented with 1 µM of 5-Ph-IAA (MedChemExpress HY-134653).

### Immunoblotting

For western-blot analysis, cells were grown under the same conditions as for the yeast growth assays. *C. albicans* cells were harvested after 5 h of growth in SD or SD+Dox medium. *S. cerevisiae* cells were harvested after 15 min of growth in SD-ura or SD-ura containing 5-Ph-IAA. Cells were pelleted and flash-frozen. Cell pellets were thawed in TENG-300 Buffer (50 mM Tris-HCl pH 7.4, 300 mM NaCl, 1 mM EDTA pH 8.0, 0.5% NP-40, 10% glycerol) supplemented with 5 mM 2-mercaptoethanol) and cOmplete ULTRA EDTA-free protease inhibitor cocktail (Roche 06538282001, 1 tablet for 50 mL of buffer). Lysis was performed under cold conditions using 0.7 mm diameter zirconia beads in a FastPrep-24 5G homogeniser (MP Biomedicals) for 2 cycles of 45 seconds at 6.5 m/s. Lysates were cleared by centrifugation, and the total protein concentration was determined using a Bradford assay. Proteins were denatured by mixing the lysates with NuPAGE LDS Sample Buffer and Reducing Agent (Invitrogen), each diluted to 1x, followed by heating for 10 min at 70 °C. Ten µg of proteins were loaded on a 3-8% Tris-acetate NuPAGE gel, run in 1x NuPAGE Tris-Acetate Running Buffer (Invitrogen) and transferred using a Trans-Blot Turbo system (Bio-Rad). Membranes were probed with the following antibodies: anti-v5 (Invitrogen R960-25), anti-Flag M2 (Sigma F3165), anti-HA (Roche 11867423001), anti-PGK1 22C5D8 (ThermoFisher 459250), anti-mouse IgG HRP (ThermoFisher 31430) and anti-rat (Dako P0450). Membranes were imaged using a Fusion FX (Vilber).

### Recombinant proteins expression and purification

For crystallisation, the DNA region encoding *C. albicans* Rcs1 was PCR amplified from genomic DNA and sub-cloned into pCR-bluntII-TOPO (Invitrogen). The fragment encoding *C. albicans* Rsc1-BD2 (residues 225-340), was then PCR amplified and cloned into a pETM11 vector as a fusion construct bearing an N-terminal His tag followed by a TEV protease cleavage site.

For native protein expression, transformed *E. coli* BL21(DE3) cells (New England Biolabs C2527I) were grown at 37 °C in LB medium containing 50 μg/mL kanamycin until reching an OD_595nm_ of 0.5. Protein expression was induced with 0.5 mM IPTG, and cultures were further incubated at 16 °C for ∼16 h.

For selenomethionine-labeled protein expression, transformed *E. coli* BL21(DE3) cells were grown at 37 °C in M9 medium (10 g/L Na_2_HPO_4_-H_2_O, 3 g/L KH_2_PO_4_, 0.5 g/L NaCl, 1 g/L NH_4_Cl) containing 2% glucose, 2 mg/L thiamine, 1 mM MgSO_4_, 0.1 mM CaCl_2_ and 50 μg/mL kanamycin, until reaching an OD_595nm_ of 0.5. The medium was supplemented with 100 mg/L each of lysine, phenylalanine and threonine and 50 mg/ml each of isoleucine, leucine, and selenomethionine. Expression was induced with 0.5 mM IPTG, and cultures were further incubated at 16 °C for ∼16 h. For both native and selenomethionine-labeled proteins, cells were collected by centrifugation, flash-frozen in liquid nitrogen and stored at -80 °C.

After thawing, the samples were handled in cold conditions. Cells were resuspended in Lysis Buffer (50 mM Tris-HCl pH 7.5, 300 mM NaCl, 10% glycerol, 15 mM imidazole, 5 mM 2-mercaptoethanol) supplemented with 1 mM Phenylmethanesulfonyl fluoride (PMSF, Thermo Scientific 215740100) and protease inhibitor cocktail (Roche 06538282001) and were lysed by sonication. The cleared lysate was incubated with Ni-NTA resin (Macherey-Nagel 745400.25) and sequentially washed with Lysis Buffer, Wash Buffer 1 (50 mM Tris-HCl pH 7.5, 500 mM NaCl, 15 mM imidazole, 5 mM 2-mercaptoethanol) and Wash Buffer 2 (as Wash Buffer 1 but with 150 mM NaCl). Proteins were eluted in Elution Buffer (same as Wash Buffer 2 but with 300 mM imidazole), dialysed overnight in the presence of His-tagged TEV protease against Dialysis Buffer (50 mM Tris-HCl pH 7.5, 150 mM NaCl, 1 mM dithiothreitol [DTT]). After dialysis, Ni-NTA resin was used to remove His-tagged species. Proteins were further purified on a Superdex 75 16/60 (GE Healthcare) column in 25 mM Hepes pH 7.5, 150 mM NaCl, 0.5 mM DTT. Proteins were concentrated to ∼60 mg/mL on a Centricon device (Millipore).

For EMSA experiments, DNA fragments encoding WT and mutated *C. albicans* Rsc1-BD2 residues 213-599, *S. cerevisiae* Rsc1-BD2 residues 221-596 and *S. cerevisiae* Rsc2-BD2 residues 261-636 were cloned into a pETM11 vector. *S. cerevisiae* constructs included an additional C-terminal 3xFlag tag.

Proteins were expressed as described above and purified via a single Ni-NTA affinity step. They were then dialysed and concentrated to at least 4 mg/mL using the purification buffers specified above.

### Protein crystallisation and structure determination

Selenomethionine-labelled Rsc1-BD2 (residues 225-340) was crystallised by the sitting drop vapor diffusion method at 4 °C using a Cartesian PixSys 4200 crystallisation robot at the High Throughput Crystallisation Laboratory of the EMBL Grenoble (https://htxlab.embl.org). Crystals were obtained by mixing 100 nL of protein (35 mg/mL) with 100 nL of reservoir solution containing 0.1 M potassium phosphate monobasic/sodium phosphate dibasic (pH 6.2) and 2-methyl-2,4-pentanediol 35% (v/v). Crystals were harvested directly from the crystallisation drop and flash-cooled in liquid nitrogen. X-ray diffraction data were collected at ESRF beamline ID30A-1 and processed using using XDS (71) and programs of the CCP4 suite (72). Substructure determination and phase calculation were performed with SHELXD and SHELXE, respectively (73). The initial electron density map was partially autotraced with SHELXE, followed by further manual model building in Coot (74) and refinement with Phenix (75). Structural figures were generated using PyMOL (Schrödinger LLC. https://www.pymol.org/pymol). Refinement and structural analysis software was compiled by SBGrid (76). Data collection and refinement statistics are summarised in **Supplementary Table 1**.

### Biophysical characterisation of recombinant proteins

Analytical gel filtration was performed by injecting 250 µg of protein on an S200 5/150 GL increase column (Cytiva) in 50 mM Tris-HCl pH 7.5, 150 mM NaCl and 1 mM DTT. Differential scanning fluorimetry (DSF) (77) was performed on a CFX96 touch real-time PCR detection system (Bio-Rad) using 0.2 mL 8-tube PCR strips (Bio-Rad TB0201) sealed with optical flat 8-cap strips (Bio-Rad TCS0803). The fluorescence of samples containing 1.5 and 3 mg/mL of protein and 1× SYPRO Orange dye (Molecular Probes) was monitored as the temperature was gradually increased from 20 °C to 100 °C at a rate of 0.5 °C/min. The wavelengths used for SYPRO Orange excitation and emission were 470 and 570 nm, respectively. The melting temperature (T_m_) was determined from the first derivative of the fluorescence melting curves using CFX Maestro software. Protein thermal stability was also assessed by nano-DSF. The intrinsic tryptophan fluorescence of proteins (0.5 to 4 mg/mL) was monitored at 330 nm and 350 nm over a temperature range of 20 °C to 90 °C with a 1 °C/min increasing temperature ramp using a Prometheus NT.48 (NanoTemper) and standard grade capillaries (Nanotemper PR-C002). T_m_ values were determined at the inflection points of the first derivative of the tryptophan fluorescence ratio (F350 nm/F330 nm) curves.

### Electrophoretic Mobility Shift Assays (EMSAs)

601 DNA probes of 197 and 145 bp were obtained by EcoRV digestion of plasmids containing repeats of the probe sequences (78). Probes were separated from the plasmid backbone by differential precipitation with polyethylene glycol 6000 and NaCl (final concentrations of 6.5% and 0.5 M, respectively) for 1 h on ice. Following centrifugation, the probes were recovered from the supernatant by ethanol-precipitation and resuspended in TE Buffer.

Oligonucleotide probes were prepared by annealing complementary single-stranded oligonucleotides (**Supplementary Table 4**) in Annealing Buffer (10 mM Tris-HCl pH 7.5, 150 mM NaCl, 1 mM EDTA). The resulting double-stranded DNA with a protruding guanine was fluorescently labelled by end-filling, incubating 4 pmol of DNA with 1 unit of Klenow fragment polymerase and 8 pmol of Cy5-dCTP (Cytiva PA55021) in Klenow Buffer for 1 h at 37 °C, followed by enzyme inactivation at 65 °C for 15 min.

Binding reactions were performed in a 10 µL reaction mix containing 50 mM Tris-HCl pH 7.5, 100 mM NaCl, 2.5% glycerol, 0.1 mM EDTA and 1 mM DTT. Variable concentrations of proteins were incubated for 30 min on ice, with 150 nM of 601 DNA probe, or with 20 nM of fluorescently labelled DNA probes and 28 ng/mL of salmon sperm DNA as nonspecific unlabelled competitor. The binding reactions were resolved on a 6% acrylamide TBE native gel (pre-run) in 0.5x TBE Buffer at 90 V at 4°C, for 70 to 90 min. Gels with the 601 DNA probes were stained for 15 min in a SYBR Safe solution (Invitrogen S33102). Gels were imaged using a Fusion FX (Vilber) and signal quantification was performed using ImageJ.

### Purification of endogenous RSC complex from *C. albicans*

*RSC Flag-immunoprecipitation for mass spectrometry. C. albicans* cells were grown in YPD medium to an OD_595nm_ of 0.8, harvested by centrifugation, flash-frozen in liquid nitrogen and lysed by cryo-milling (Retsch MM 400, 2 cycles of 1 min at 30 Hz). All subsequent steps were conducted at 4°C. The resulting cell powder was resuspended in Resuspension Buffer A (50 mM Tris-HCl pH 7.5, 150 mM NaCl, 0.5 mM EDTA, 0.5 % NP 40, 1 mM DTT, 1 mM PMSF, protease inhibitor cocktail) and cleared by centrifugation. The cleared lysate was incubated with magnetic anti-Flag beads (MCE HY-K0207) for 2 h. Beads were washed using Wash Buffer A (identical to Resuspension Buffer A but with 500 mM NaCl), incubated for 1 h with Benzonase Buffer (50 mM Tris-HCl pH 7.5, 150 mM NaCl, 1 mM Mg acetate, 0.5 % NP-40, 1 mM DTT) containing 40 U/mL of Benzonase nuclease (Sigma-Aldrich E1014), and washed using Wash Buffer A. The RSC complex was eluted in Resuspension Buffer A supplemented with 1 mg/mL of 3xFlag peptide (MCE HY-P0319). The eluate was mixed with NuPAGE LDS Sample Buffer and Reducing Agent (Invitrogen), each diluted to 1×, and heat-denatured for 10 min at 70 °C. RSC purified complexes were visualised by SDS-PAGE followed by silver staining using the SilverQuest kit (Invitrogen LC6070).

*RSC TAP for REA assay. C. albicans* cells were grown in YPD medium to stationary phase, harvested and lysed as described for Flag-immunoprecipitation. The cell powder obtained from 8 L of culture (∼300 g) was resuspended in Resuspension Buffer B (50 mM Tris-HCl pH 7.5, 100 mM (NH4)_2_SO_4_, 10 % glycerol, 0.5 mM EDTA, 2 mM MgCl_2_, 0.5 % NP-40, 1 mM PMSF, protease inhibitor cocktail) supplemented with 7.5 kU of Benzonase nuclease, incubated for 40 min and cleared by centrifugation. The cleared lysate was incubated for 6 h with IgG Sepharose 6 Fast Flow beads (Cytiva 17096901). Beads were washed with TEV Cleavage Buffer (50 mM Tris-HCl pH 7.5, 150 mM NaCl, 10 % glycerol, 0.5 mM EDTA, 3 mM CaCl_2_, 1 mM DTT, 1 mM PMSF) and incubated for 16 h with Calmodulin Sepharose 4B beads (Cytiva 17052901) and 200 µg of TEV protease. Beads were successively washed with Wash Buffer B (50 mM Tris-HCl pH 7.5, 150 mM NaCl, 10% glycerol, 0.5 mM EDTA, 2 mM CaCl_2_, 1 mM DTT, 1 mM PMSF) and Wash Buffer B containing 400 mM NaCl. The complex was eluted in Elution Buffer B (50 mM Tris-HCl pH 7.5, 150 mM NaCl, 10% glycerol, 0.5 mM EDTA, 2 mM EGTA, 1 mM DTT, 1 mM PMSF) and concentrated using a Vivaspin concentrator column (Cytiva 28932363). RSC purified complexes were visualised by silver stained SDS-PAGE. The relative concentrations of RSC complexes, purified from *C. albicans* strains expressing WT or mutant TAP-tagged Rsc1 subunit, were determined by western-blot quantification of Rsc1 using an anti-TAP tag antibody (Invitrogen CAB1001).

### Mass-spectrometry-based quantitative proteomic analysis

Eluted proteins solubilized in LDS buffer were stacked in the top of a NuPAGE 4-12 % gel (Invitrogen) and stained with Coomassie Blue R-250 (Bio-Rad) before in-gel digestion using modified trypsin (Promega, sequencing grade) as previously described (79). The resulting peptides were analysed by online nano-liquid chromatography coupled to tandem MS (Ultimate 3000 and Q-Exactive HF, Thermo Scientific). Peptides were sampled on a PepMap C18 precolumn (300 µm x 5 mm, Thermo Scientific) and separated on a 75 µm x 250 mm C18 column (Aurora Generation 3, 1.7μm, IonOpticks) using a 120-min acetonitrile gradient. MS and MS/MS data were acquired using Xcalibur (Thermo Scientific).

Peptides and proteins were identified by Mascot (version 2.8.0, Matrix Science) through concomitant searches against the Uniprot database (*Candida albicans* SC5314 taxonomy, 20220527 download) and a homemade database containing the sequences of classical contaminant proteins found in proteomic analyses (keratins, trypsin, etc.). Trypsin/P was chosen as the enzyme and two missed cleavages were allowed. Precursor and fragment mass error tolerances were set at respectively at 10 and 20 ppm. Peptide modifications allowed during the search were: Carbamidomethyl (C, fixed), Acetyl (Protein N-term, variable) and Oxidation (M, variable). The Proline software (version 2.2) (80) was used for the compilation, grouping and filtering of the results (conservation of rank 1 peptides, peptide length ≥ 6 amino acids, false discovery rate of peptide-spectrum-match identifications < 1% (81), and minimum of one specific peptide per identified protein group). Proline was then used to perform a MS1-based label-free quantification of the identified protein groups based on specific and razor peptides, with the cross-assignment option enabled. MS data have been deposited to the ProteomeXchange Consortium via the PRIDE partner repository (82) with the dataset identifier PXD062102.

Statistical analyses were performed using the Prostar software (version 1.34.5) (83). After log2 transformation, extracted abundance values were median normalized before missing value imputation (SLSA algorithm for partially observed values in the condition and DetQuantile algorithm for totally absent values in the condition). Statistical testing was conducted using limma, whereby differentially expressed proteins were selected using a log2(Fold Change) cut-off of 2 and a p-value cut-off of 0.01, allowing to reach a false discovery rate inferior to 1% according to the Benjamini-Hochberg estimator. Proteins found differentially abundant but identified by MS/MS in less than one replicate or with at least one imputed value in the condition in which they were found to be more abundant were manually invalidated (p-value = 1).

Stoichiometries of proteins found enriched with Flag-tagged Rsc1 were estimated using intensity-based absolute quantification (iBAQ) (84) values extracted by Proline software, as described in (85).

### Restriction Enzyme Accessibility (REA) assay

REA assays were performed in an 8 µL reaction volume containing 20 mM Tris-HCl pH 7.5, 50 mM NaCl, 10 mM MgCl_2_, 0.5 % Tween 20, 0.1 mg/mL bovine serum albumin and 1 mM DTT. Remodeling reactions containing 20 nM of mononucleosome (EpiDyne ST601-GATC, EpiCypher 6-4111), 2 µL of TAP-purified RSC complex, 1 mM of ATP (Thermo Scientific R1441) and 20 U of DpnII (New England Biolabs R0543L) were incubated for 8 h at 30 °C. Reactions were stopped by the addition of 8 µL of Quench Buffer (10 mM Tris pH 7.5, 40 mM EDTA, 0.6% SDS) supplemented with 50 μg/mL of Proteinase K (Invitrogen 11588916), followed by incubation for 20 min at 55 °C. The resulting DNA samples were migrated on 8% native polyacrylamide gels (prepared and run in 0.5× TBE). Gels were stained for 15 min in a SYBR Safe solution and imaged using a Fusion FX (Vilber).

### Micrococcal nuclease digestion and high-throughput sequencing (MNase-seq)

*C. albicans* cells were grown overnight at 30 °C in Synthetic Defined (SD) yeast medium with agitation. Cultures were diluted in SD or in SD+Dox at an OD_595nm_ of 0.1 and grown for 5 h. Cells were crosslinked with 1% formaldehyde for 10 min, at room temperature. Crosslinking reaction was quenched by addition of glycine to a final concentration of 150 mM. Cells were pelleted by centrifugation at 4 °C, washed with cold PBS and pellets were flash frozen and stored at -80 °C. Pellets originating from 25 mL of culture were thawed in 0.5 mL of Spheroplast Buffer (1 M Sorbitol, 50 mM Tris-HCl pH 7.5, 5 mM 2-mercaptoethanol) and treated with 0.75 mg of zymolyase (Roth 9329.1) for 10 min at room temperature. Spheroplasts were washed once with Spheroplast Buffer and once with MNase Digestion Buffer (1 M Sorbitol, 10 mM Tris-HCl pH 7.5, 50 mM NaCl, 5 mM MgCl_2_, 1 mM CaCl_2_, 0.075% NP-40, 0.5 mM spermidine, 1 mM 2-mercaptoethanol). For chromatin digestion, spheroplasts were resuspended in 50 µL of MNase Digestion Buffer and treated with 3U of MNase (Thermo Scientific EN0181) for 45 min at 37 °C with agitation, for chromatin digestion. Digestion was stopped by the addition of EDTA and EGTA to a final concentration of 5 mM each. Samples were incubated with 20 µg of RNase A for 20 min at 37 °C. Crosslinks were reversed by the addition of 1 volume of Decrosslinking Buffer (10 mM Tris-HCl pH 7.5, 50 mM NaCl, 0.075 % NP-40, 2 % SDS), 100 µg of Proteinase K (Invitrogen 11588916) and incubation at 62 °C for 4 h with agitation. DNA was recovered by two cycles of phenol:chloroform:isoamyl alcohol (25:24:1) extraction, ethanol precipitated, resuspended in 20 µL of TE Buffer (10 mM Tris-HCl pH 8, 1 mM EDTA) and treated with 20 µg of RNase A for 45 min at 37 °C. 50 ng of DNA were used for library preparation, using the KAPA Hyper Prep kit, Universal Adapter and UDI Primer Mixes (Roche 7962347001, 9063781001 and 9134336001, respectively), following the supplier’s recommendations. For each sample, the number of PCR cycles required for library amplification was determined by qPCR, ensuring amplification occurred within the exponential phase. Library size distributions and concentrations were analysed using a Bioanalyzer 2100 (Agilent) and a Qubit fluorometer (Thermo Scientific). Pooled libraries were sequenced on a HiSeq X (Illumina) using paired-end mode.

### MNase-seq data analysis

To process MNase-seq raw data, demultiplexed fastq files were pre-processed using Cutadapt (86) for adapter removal and reads trimming (parameters --quality-cutoff 33 --length 30 --minimum-length 20). Trimmed paired reads were aligned to a haploid version of *C. albicans* genome (SC5314, Assembly 22) using Bowtie2 (87) (parameters --end-to-end --no-mixed --no-discordant). Aligned reads were filtered for mapping quality (q 30) using SAMtools (88). The coverage was calculated using deepTools bamCoverage (89), with single-base resolution (--binSize 1), by extending reads to match the fragment size defined by the two read mates (--extendReads), normalising by Reads Per Kilobase per Million mapped reads (--normalizeUsing RPKM), filtering for fragments in the 100–200 bp range (--minFragmentLength 100 --maxFragmentLength 200), and by centring reads with respect to the fragment length (--centerReads). *C. albicans* genes annotations (from TSS to TES) were downloaded from https://github.com/tschemic/ATACseq_analysis (90). The nucleosome occupancy (or total coverage) over protein coding genes was calculated using deepTools computeMatrix and heatmaps were plotted using deepTools plotHeatmap (89).

To analyse the +1 nucleosome shifts, MNase-seq BAM files were processed using the nucleR package (–p1.max.dowstream 30 –window 500) (91) for detection of nucleosome peaks adjacent to the TSS of protein coding genes. The obtained nucleosome positions were crossed with gene annotations to ensure that were attributed to genuine TSS and not TES. For each individual gene, the distance between the TSS and the +1 nucleosome was computed, distances exceeding 150 bp were considered as non-reliable and filtered out. For each gene, the shift of the +1 nucleosome upon Dox addition was calculated as follow: the position of the +1 in a -Dox sample was compared to its position in the matched +Dox sample; only genes where a +1 nucleosome peak was detected in both matched ±Dox samples were taken into account.

To relate our MNase-seq data to gene expression levels, we used publicly available RNA-seq data as a proxy of gene expression. The RNA-seq data were originally obtained in *C. albicans* WT cells during exponential growth in five replicates (GSM1893022, GSM1893023, GSM1893024, GSM1893025, GSM1893026) (47). We used these data reprocessed by (48), extracted FPKM values and averaged the replicates. We stratified genes into quartiles of gene expression based on the mean FPKM value.

To perform a *de novo* motif search in the C*. albicans* NDR, we extracted sequences from the interval -300 to +100 bp around the TSS of all protein-coding genes in the *C. albicans* genome. The sequences were used as input to run the MEME suite (92) (-minw 6 -maxw 15 and forcing all sequences to be analysed). To reduce computational time, the sequences were randomly split into 3 sets before running MEME. All 3 groups gave very similar results, the motif from one group was used as representative. The motif quality was evaluated using ROC area under the curve, where the positive regions were the interval -300 to +100 bp around the TSS, while negative regions were -800 to -400 and +200 to +500 bp intervals relative to TSS. The matrix of the first set of genes was used to perform a genomic distribution of scores. The top 99.5% best score was used as the threshold for good quality binding site. Using this threshold and matrix, putative binding sites were identified in the genome.

To analyse nucleosome positioning and gene expression in *S. cerevisiae*, we used published datasets: GSE69400/GSM1700669_WT_A for MNase-seq (93), and Rpo21 (RNAPII) CRAC signal over protein coding genes (GSE69676) was used as a proxy of gene expression (94).

### ampDAP-seq

The coding sequence of *S. cerevisiae* Rsc2 was fused to three repeats of the Flag epitope tag and cloned in the pTNT vector (Promega). We used the ampDAP-seq libraries described in (95). ampDAP-seq experiments were performed following a previously described protocol (49). The analysis of ampDAP-seq sequences (peak finding, motif search, and evaluation) was performed as described in (96). 50µL of TNT reaction was incubated for 1 h at 4 °C on a rotating wheel. Beads were then immobilised and washed 6 times with 100 µL of DAP buffer (PBS 1x, 5 mM TCEP, 0.0005 % NP-40), moved to a new tube and washed once again. DAP-seq input libraries (50 ng) were then added, and protein-DNA mixes were incubated for 1.5 h at 4 °C on a rotating wheel. Beads were immobilised and washed 5 times with 100 µl DAP buffer, moved to a new tube and washed twice. Finally, beads were mixed with 30 µl of elution buffer (10 mM Tris-HCl pH 8.5) and heated for 10 min at 90 °C. IP-ed DNA fragments contained in the elution were amplified by PCR according to published protocol (49) with Illumina TruSeq primers. PCR products were purified using AMPure XP magnetic beads (Beckman Coulter) following manufacturer’s instructions. Library molar concentrations were determined by qPCR using NEBNext Library Quant Kit for Illumina (NEB). Libraries were then pooled with equal molarity and sequenced using Illumina HiSeq. This experiment was performed in duplicate.

The analysis of ampDAP-seq sequences (peak finding, motif search, and evaluation) was performed as described in (96). Briefly, each replicate was mapped to the S. cerevisiae (S288C_reference_sequence_R64-3-1) using Bowtie2 (87) with default parameters. After filtering low quality mapped reads and PCR duplicates, peaks were called using MACS2 (in paired-end mode, with –call-summit parameter). Peaks from each duplicate were compiled using MPSC to obtain robust consensus peaks. *De novo* motif search was performed on all consensus peaks using MEME suite (-minw 4 -maxw 20).

### Protein sequence analysis and structural prediction

Three-dimensional structure images were done using PyMOL (www.pymol.org), sequence alignment was calculated using Clustal Omega (97) and visualised using ESPrit3.0 (https://espript.ibcp.fr/ESPript/cgi-bin/ESPript.cgi). Surface conservation was computed using the Consurf server (https://consurf.tau.ac.il/consurf_index.php). The structural prediction of *C. albicans* Rsc1, residues 213-599, was obtained using AlphaFold2 (40).

### Phylogenetic analysis of fungal BDs

The evolutionary conservation of the functional determinants of Rsc1-BD2 was investigated using 15 fungal species, listed in **Table S5**. These species were selected based on their representativeness of the fungal kingdom, as well as the availability of reliable and well-annotated proteome data. For each species, we identified all proteins containing at least one BD (**Table S5**). This approach allowed us to extract 209 BD sequences, which were aligned using the Clustal Omega tool (97). Residues corresponding to the structural determinants of Rsc1-BD2 DNA-binding surface (basic residues located on alpha-Z helix), as well as the acetylated lysine-binding pocket (Y and N residues required for coordinating acetyl group binding), were analysed and their conservation was visualised as a heatmap.

## Data availability

The crystal structure and diffraction data for Rsc1 BD2 have been deposited in the Protein Data Bank under accession code 9QHR. MNase-seq sequences generated during this work will be deposited in the GEO database (http://www.ncbi.nlm.nih.gov/geo/) under accession numbers XXXXXXXX. Mass spectrometry data have been deposited to the ProteomeXchange Consortium via the PRIDE partner repository with the dataset identifier PXD062102.

## Acknowledgments

This work was supported by a fellowship from the Fondation pour la Recherche Médicale (FRM grant ARF201809007076) to CS, by grants from the Agence Nationale de Recherche (ANR-18-CE18-0007, ANR-21-CE18-0041) to JG and CP and by a GRAL PhD fellowship to RD. This work used the platforms of the Grenoble Instruct-ERIC center (ISBG; UAR 3518 CNRS-CEA-UGA-EMBL) within the Grenoble Partnership for Structural Biology (PSB), supported by FRISBI (ANR-10-INBS-0005-02) and GRAL, financed within the University Grenoble Alpes Graduate School (Ecoles Universitaires de Recherche) CBH-EUR-GS (ANR-17-EURE-0003). We acknowledge the European Synchrotron Radiation Facility (ESRF) for provision of synchrotron radiation facilities under proposal number MX-2329 and thank Didier Nurizzo and Matthew Bowler for assistance and support in using beamline ID30A-1. The proteomic experiments were partially supported by Agence Nationale de la Recherche under projects ProFI (Proteomics French Infrastructure, ANR-10-INBS-08) and GRAL (ANR-17-EURE-0003). The authors thank Carlos Fernandez-Tornero and members of the Petosa group for helpful advice in protein biochemistry, members of the Govin group for their support in the lab and for fostering a collaborative working environment, and Charles McKenna for insightful discussions on chemical inhibition.

## Author contributions

CS, JG, CP performed conceptualisation; CS, JG, CP, FP performed data curation; CS, JLu, JG, CP, YC performed formal analysis; JG, CS, CP performed funding acquisition; CS, OV, JLy, RD, ET, JM, AA performed investigation; CS, JG, CP, YC, FP performed methodology; CS, JG performed project administration; CS, OV, JLy, JLu, RD, ET, JM, AA, YC, FP, CP, JG provided resources; CS, JG, CP, FP, YC performed supervision; CS, JG, CP, FP, YC performed validation; CS, JG performed visualization; CS wrote the original draft; JG, CP, YC, CS wrote, reviewed and edited.

