## Supplementary_materials for "Rsc1 DNA-binding bromodomain drives RSC activity at A-rich promoters"

#### Supplementary table 1. Crystallographic data collection and refinement statistics

| | PDB 9QHR |
|---|---|
| **Data Collection**[1] | |
| Synchrotron beamline | ESRF ID30A-1 |
| Wavelength (Å) | 0.9655 |
| Space group | $P2_12_12_1$ |
| Unit cell dimensions | |
| *a* (Å) | 66.18 |
| *b* (Å) | 68.65 |
| *c* (Å) | 130.27 |
| Molecules in asym. unit | 4 |
| Resolution range (Å) | 38.5 – 1.50 |
| (outer shell) | (1.61– 1.50) |
| No. of measured reflections | 1,095,529 (113,079) |
| No. of unique reflections | 181,652 (31,375) |
| Multiplicity | 6.03 (3.60) |
| Completeness (%) | 99.0 (94.7) |
| Mean I/sigma(I) | 14.1 (1.2) |
| $R_{meas}$ | 0.072 (1.15) |
| $CC_{1/2}$ | 0.999 (0.628) |
| **Refinement** | |
| Resolution used for refinement | 38.5 – 1.50 |
| Reflections used (total/$R_{free}$) | 181,652 / 1900 |
| $R_{work}$/$R_{free}$ | 0.2018 / 0.2261 |
| No. of atoms/<B-factor> (Å$^2$) | |
| Protein | 3892 / 28.6 |
| Water | 722 / 36.0 |
| RMS deviations: | |
| Bond distances (Å) | 0.006 |
| Bond angles (º) | 0.744 |
| Ramachandran analysis (%) | |
| Favored/ outliers | 99.8 / 0.0 |
| Molprobity analysis | |
| Clash Score / Overall score | 2.43 / 1.03 |

[1] Numbers in parentheses refer to the outer resolution shell.

### Supplementary Table 2. Yeast strains

| Name | ID | Species | Genotype information | | | | Reference |
|---|---|---|---|---|---|---|---|
| | | | ***RSC1* locus** | | | | |
| WT (SN152)<br>*ura3::imm434::URA3/ura3::imm434 iro1::IRO1/iro1::imm434 his1::hisG/his1::hisG leu2/leu2 arg4/arg4* | yCaJG03 | *C. albicans* | *RSC1//RSC1* | | | | Noble and Johnson, 2005 |
| pTet::RSC1//rsc1Δ | yCaJG143 | *C. albicans* | *ARG4::tetO-pTet::3v5-RSC1//rsc1Δ::NatMX6* | | | | This study |
| pTet::RSC1//RSC1-WT | yCaJG144 | *C. albicans* | *ARG4::tetO-pTet::3v5-RSC1//RSC1-3xFlag::LEU2* | | | | This study |
| pTet::RSC1//rsc1-Δbd1 | yCaJG145 | *C. albicans* | *ARG4::tetO-pTet::3v5-RSC1//rsc1-Δbd1-3xFlag::LEU2* | | | | This study |
| pTet::RSC1//rsc1-Δbd2 | yCaJG146 | *C. albicans* | *ARG4::tetO-pTet::3v5-RSC1//rsc1-Δbd2-3xFlag::LEU2* | | | | This study |
| pTet::RSC1//rsc1-bd1Y45F | yCaJG147 | *C. albicans* | *ARG4::tetO-pTet::3v5-RSC1//rsc1-bd1Y45F-3xFlag::LEU2* | | | | This study |
| pTet::RSC1//rsc1-bd2Y267F | yCaJG148 | *C. albicans* | *ARG4::tetO-pTet::3v5-RSC1//rsc1-bd2Y267F-3xFlag::LEU2* | | | | This study |
| pTet::RSC1//rsc1-Δbd1-Δbd2 | yCaJG149 | *C. albicans* | *ARG4::tetO-pTet::3v5-RSC1//rsc1-Δbd1-Δbd2-3xFlag::LEU2* | | | | This study |
| pTet::RSC1//rsc1-bd1Y45F-bd2Y267F | yCaJG150 | *C. albicans* | *ARG4::tetO-pTet::3v5-RSC1//rsc1-bd1Y45F-bd2Y267F-3xFlag::LEU2* | | | | This study |
| pTet::RSC1//rsc1-bd2Y267A-Y310A | yCaJG153 | *C. albicans* | *ARG4::tetO-pTet::3v5-RSC1//rsc1-bd2Y267A-Y310A-3xFlag::LEU2* | | | | This study |
| pTet::RSC1//rsc1-bd2Y267R-Y310R | yCaJG154 | *C. albicans* | *ARG4::tetO-pTet::3v5-RSC1//rsc1-bd2Y267R-Y310R-3xFlag::LEU2* | | | | This study |
| pTet::RSC1//rsc1-bd2Y267A-Y310R | yCaJG155 | *C. albicans* | *ARG4::tetO-pTet::3v5-RSC1//rsc1-bd2Y267A-Y310R-3xFlag::LEU2* | | | | This study |
| pTet::RSC1//rsc1-bd2-K238A-K241A-K242A | yCaJG177 | *C. albicans* | *ARG4::tetO-pTet::3v5-RSC1//RSC1-bd2-K238A-K241A-K242A-3xFlag::LEU2* | | | | This study |
| pTet::RSC1//rsc1-bd2-K238E-K241E-K242E | yCaJG178 | *C. albicans* | *ARG4::tetO-pTet::3v5-RSC1//RSC1-bd2-K238E-K241E-K242E-3xFlag::LEU2* | | | | This study |
| pTet::RSC1//rsc1-athook-R218A-R219A-R221A | yCaJG179 | *C. albicans* | *ARG4::tetO-pTet::3v5-RSC1//RSC1-athook-R218A-R219A-R221A-3xFlag::LEU2* | | | | This study |
| pTet::RSC1//rsc1-athook-R221A | yCaJG180 | *C. albicans* | *ARG4::tetO-pTet::3v5-RSC1//RSC1-athook-R221A-3xFlag::LEU2* | | | | This study |
| pTet::RSC1//rsc1-athook-R221E | yCaJG181 | *C. albicans* | *ARG4::tetO-pTet::3v5-RSC1//RSC1-athook-R221E-3xFlag::LEU2* | | | | This study |
| pTet::RSC1//rsc1-bah-R477A-R487A-R489A | yCaJG182 | *C. albicans* | *ARG4::tetO-pTet::3v5-RSC1//RSC1-bah-R477A-R487A-R489A-3xFlag::LEU2* | | | | This study |
| pTet::RSC1//rsc1-bah-R477E-R487E-R489E | yCaJG183 | *C. albicans* | *ARG4::tetO-pTet::3v5-RSC1//RSC1-bah-R477E-R487E-R489E-3xFlag::LEU2* | | | | This study |
| RSC1-TAP | yCaJG151 | *C. albicans* | *RSC1-TAP::ARG4//RSC1* | | | | This study |
| rsc1-bd2-K238E-K241E-K242E-TAP | yCaJG184 | *C. albicans* | *rsc1-bd2-K238E-K241E-K242E-TAP::ARG4//RSC1* | | | | This study |
| | | | ***RSC1* locus** | ***RSC2* locus** | **Other genomic modifications** | **Plasmid** | |
| WT (BY4741)<br>*MATa; his3Δ1; leu2Δ0; met15Δ0; ura3Δ0* | yJG707 | *S. cerevisiae* | *RSC1* | *RSC2* | *his3Δ1* | | Brachmann et al., 1998 |
| AID-RSC1 AID-RSC2 | yJG754 | *S. cerevisiae* | *3xAID-3xv5-RSC1* | *3xAID-3xHA-RSC2* | *OsTIR1-F74A::NatMX6::his3Δ1* | | This study |
| AID-RSC1 AID-RSC2 pRS316::RSC2-WT | yJG756 | *S. cerevisiae* | *3xAID-3xv5-RSC1* | *3xAID-3xHA-RSC2* | *OsTIR1-F74A::NatMX6::his3Δ1* | *pRS316::3xFlag-RSC2* | This study |
| AID-RSC1 AID-RSC2 pRS316::rsc2-athook-R269E | yJG757 | *S. cerevisiae* | *3xAID-3xv5-RSC1* | *3xAID-3xHA-RSC2* | *OsTIR1-F74A::NatMX6::his3Δ1* | *pRS316::3xFlag-rsc2-athook-R269E* | This study |
| AID-RSC1 AID-RSC2 pRS316::rsc2-bah-R514E-K524E-R526E | yJG758 | *S. cerevisiae* | *3xAID-3xv5-RSC1* | *3xAID-3xHA-RSC2* | *OsTIR1-F74A::NatMX6::his3Δ1* | *pRS316::3xFlag-rsc2-bah-R514E-K524E-R526E* | This study |
| AID-RSC1 AID-RSC2 pRS316::rsc2-bd2-K286E-K289E-K290E | yJG759 | *S. cerevisiae* | *3xAID-3xv5-RSC1* | *3xAID-3xHA-RSC2* | *OsTIR1-F74A::NatMX6::his3Δ1* | *pRS316::3xFlag-rsc2-bd2-K286E-K289E-K290E* | This study |
| AID-RSC1 AID-RSC2 pRS316 | yJG764 | *S. cerevisiae* | *3xAID-3xv5-RSC1* | *3xAID-3xHA-RSC2* | *OsTIR1-F74A::NatMX6::his3Δ1* | *pRS316* | This study |

### Supplementary Table 3. Plasmids

| ID | Description | Purpose | Destination species | Reference |
|---|---|---|---|---|
| pJG453 | ARG4::pTetO::3v5-RSC1 | Conditional depletion of *RSC1* | *C. albicans* | This study |
| pJG478 | rsc1Δ::NatMX6 | Deletion of *RSC1* ORF | *C. albicans* | This study |
| pJG495 | RSC1-3xFlag::LEU2 | Expression of WT Rsc1 | *C. albicans* | This study |
| pJG496 | rsc1-Δbd1-3xFlag::LEU2 | Expression of mutated Rsc1 | *C. albicans* | This study |
| pJG497 | rsc1-Δbd2-3xFlag::LEU2 | Expression of mutated Rsc1 | *C. albicans* | This study |
| pJG498 | rsc1-bd1Y45F-3xFlag::LEU2 | Expression of mutated Rsc1 | *C. albicans* | This study |
| pJG499 | rsc1-bd2Y267F-3xFlag::LEU2 | Expression of mutated Rsc1 | *C. albicans* | This study |
| pJG500 | rsc1-Δbd1-Δbd2-3xFlag::LEU2 | Expression of mutated Rsc1 | *C. albicans* | This study |
| pJG501 | rsc1-bd1Y45F-bd2Y267F-3xFlag::LEU2 | Expression of mutated Rsc1 | *C. albicans* | This study |
| pJG508 | rsc1-bd2Y267A-Y310A-3xFlag::LEU2 | Expression of mutated Rsc1 | *C. albicans* | This study |
| pJG509 | rsc1-bd2Y267R-Y310R-3xFlag::LEU2 | Expression of mutated Rsc1 | *C. albicans* | This study |
| pJG510 | rsc1-bd2Y267A-Y310R-3xFlag::LEU2 | Expression of mutated Rsc1 | *C. albicans* | This study |
| pJG535 | rsc1-bd2-K238A-K241A-K242A-3xFlag::LEU2 | Expression of mutated Rsc1 | *C. albicans* | This study |
| pJG536 | rsc1-bd2-K238E-K241E-K242E-3xFlag::LEU2 | Expression of mutated Rsc1 | *C. albicans* | This study |
| pJG537 | rsc1-athook-R218A-R219A-R221A-3xFlag::LEU2 | Expression of mutated Rsc1 | *C. albicans* | This study |
| pJG538 | rsc1-athook-R221A-3xFlag::LEU2 | Expression of mutated Rsc1 | *C. albicans* | This study |
| pJG539 | rsc1-athook-R221E-3xFlag::LEU2 | Expression of mutated Rsc1 | *C. albicans* | This study |
| pJG540 | rsc1-bah-R477A-R487A-R489A-3xFlag::LEU2 | Expression of mutated Rsc1 | *C. albicans* | This study |
| pJG541 | rsc1-bah-R477E-R487E-R489E-3xFlag::LEU2 | Expression of mutated Rsc1 | *C. albicans* | This study |
| pJG502 | RSC1-TAP::ARG4 | Purification of the RSC complex | *C. albicans* | This study |
| pJG571 | rsc1-bd2-K238E-K241E-K242E-TAP::ARG4 | Purification of the RSC complex | *C. albicans* | This study |
| pJG557 | gRNA_ScerRSC1_PAM4_in_pWS082 | CRISPR/Cas9 edifding of RSC1 | *S. cerevisiae* | This study |
| pJG558 | gRNA_ScerRSC2_PAM-26_in_pWS082 | CRISPR/Cas9 edifding of RSC2 | *S. cerevisiae* | This study |
| pJG544 | OsTIR1$^{F74A}$::NatMX6_ integrative_in_HIS3 | AID system | *S. cerevisiae* | This study |
| pRS316 | Empty vector | Control plasmid for Rsc2 expression | *S. cerevisiae* | Sikorski and Hieter, 1989 |
| pJG578 | pRS316::RSC2 | Expression of WT Rsc2 | *S. cerevisiae* | This study |
| pJG579 | pRS316::3xFlag-rsc2-athook-R269E | Expression of mutated Rsc2 | *S. cerevisiae* | This study |
| pJG580 | pRS316::3xFlag-rsc2-bah-R514E-K524E-R526E | Expression of mutated Rsc2 | *S. cerevisiae* | This study |
| pJG581 | pRS316::3xFlag-rsc2-bd2-K286E-K289E-K290E | Expression of mutated Rsc2 | *S. cerevisiae* | This study |
| pJG468 | pETM-11::Calb_Rsc1_BD2_225-340 | Production of recombinant *C. albicans* Rsc1-BD2 protein for crystallography | *E. coli* | This study |
| pJG550 | pETM11::Calb_Rsc1_213_599_WT | Production of recombinant *C. albicans* Rsc1 protein for EMSA | *E. coli* | This study |
| pJG551 | pETM11::Calb_Rsc1_213_599_athook_R221E | Production of recombinant *C. albicans* Rsc1 protein for EMSA | *E. coli* | This study |
| pJG552 | pETM11::Calb_Rsc1_213_599_bd2_K238E_K241E_K242E | Production of recombinant *C. albicans* Rsc1 protein for EMSA | *E. coli* | This study |
| pJG553 | pETM11::Calb_Rsc1_213_599_bah_R477E_R487E_R489 E | Production of recombinant *C. albicans* Rsc1 protein for EMSA | *E. coli* | This study |
| pJG554 | pETM11::Calb_Rsc1_213_599_bd2_bah_3xKE_3xRE | Production of recombinant *C. albicans* Rsc1 protein for EMSA | *E. coli* | This study |
| pJG567 | pETM11::Scer_Rsc1_221_596_WT-3xFlag | Production of recombinant *S. cerevisiae* Rsc1 protein for EMSA | *E. coli* | This study |
| pJG568 | pETM11::Scer_Rsc1_221_596_K246E_R249E_R250E-3xFlag | Production of recombinant *S. cerevisiae* Rsc1 protein for EMSA | *E. coli* | This study |
| pJG569 | pETM11::Scer_Rsc2_261_636_WT-3xFlag | Production of recombinant *S. cerevisiae* Rsc2 protein for EMSA | *E. coli* | This study |
| pJG570 | pETM11::Scer_Rsc2_261_636_K286E_K289E_K290E-3xFlag | Production of recombinant *S. cerevisiae* Rsc2 protein for EMSA | *E. coli* | This study |
| pJG565 | pTNT_3xFlag::Scer_Rsc2_261_641_WT | Cell free production of *S. cerevisiae* Rsc2 protein for ampDAP-seq | | This study |

### Supplementary Table 4. Oligonucleotides and double stranded DNA fragments

| ID | sequence | Type | Purpose | Reference |
|---|---|---|---|---|
| prJG1798 | gactGaaattatatttcaagcatgg | Oligonucleotide | CRISPR/Cas9 *S. cerevisiae*, ScerRSC1_sgRNA | This study |
| prJG1799 | aaacccatgcttgaaatataatttC | Oligonucleotide | CRISPR/Cas9 *S. cerevisiae*, ScerRSC1_sgRNA | This study |
| prJG1800 | gactGtcgcgcagaaccagacgaag | Oligonucleotide | CRISPR/Cas9 *S. cerevisiae*, ScerRSC2_sgRNA | This study |
| prJG1801 | aaaccttcgtctggttctgcgcgaC | Oligonucleotide | CRISPR/Cas9 *S. cerevisiae*, ScerRSC2_sgRNA | This study |
| gJG0060 | GAACAGTGTGGTATTAGCGAAGGGAAAATCTGTGAAGTGCCCT AGCTACAAAACAGAGATAAAAAAAATTATATTTCAAGCATGCCC AAGGATCCTGCAAAACCACCGGCTACGGCGCAGGTCGTAGGC TGGCCACCAGTTCGTTCCTACAGAAAAAATGTCATGGTTTCCT GCCAAAAGTCATCCGGAGGACCGAAGGATCCAGCAAAGCCAC CTGCGACTGCTCAGGTCGTTGGGTGGCCGCCGGTTCGTTCCT ATCGTAAGAACGTGATGGTAAGTTGTCAGAAGTCATCTGGGGG ACCGAAAGACCCGGCCAAGCCTCCTGCAACGGCGCAAGTGGT CGGCTGGCCTCCCGTGAGGAGCTATAGAAAAAATGTAATGGTA TCCTGTCAAAAATCAAGTGGGGGGTCCGGATCAGAGACGCCC GGCACTTCTGAGAGTGCAACCCCTGAAAGTGGTAAACCTATAC CGAATCCCTTGCTAGGGCTGGACTCCACCGGAGGTTCCGGGA AGCCAATTCCTAACCCTCTGTTGGGACTAGACAGCACGGGTTC TGGCGGTAAGCCTATTCCAAATCCCTTGTTGGGTCTGGACAGT ACCGGCAGTGGTGGGGGCGGATCCGTGGAGCAAGATAATGG GTTTTTACAGAAGCTACTCAAAACACAATACGATGCTGTCTTCC ATTTAAAGGATGAAAATGGTATAGAAATATATCCAATATTCAAT GTACTGCCAC | Double stranded DNA | CRISPR/Cas9 *S. cerevisiae*, repair template for 3xAID-3xv5-RSC1 | This study |
| gJG0066 | GCTGAAAAAAGTATTGGTAACAGTTCAATACGTGATCAAATATA CAGCACGTCGCGCAGAACCAGACGAAGCACAGAATATTCTAC ATTGACAGTGCATGATGCCCAAGGATCCTGCAAAACCACCGG CTACGGCGCAGGTCGTAGGCTGGCCACCAGTTCGTTCCTACA GAAAAAATGTCATGGTTTCCTGCCAAAAGTCATCCGGAGGACC GAAGGATCCAGCAAAGCCACCTGCGACTGCTCAGGTCGTTGG GTGGCCGCCGGTTCGTTCCTATCGTAAGAACGTGATGGTAAGT TGTCAGAAGTCATCTGGGGGACCGAAAGACCCGGCCAAGCCT CCTGCAACGGCGCAAGTGGTCGGCTGGCCTCCCGTGAGGAG CTATAGAAAAAATGTAATGGTATCCTGTCAAAAATCAAGTGGG GGGTCCGGATCAGAGACGCCCGGCACTTCTGAGAGTGCAACC CCTGAAAGTGCAGCGGTATACCCATATGACGTACCAGATTATG CTGGTTATCCGTACGACGTTCCCGATTATGCCGGGTCTTACCC CTATGACGTCCCTGATTACGCACCGGCGGCTGCCGCCCCTGA TGACAATTCAAACTCGTCCACTCAAAACTCAAGCGCATTATACA AGGACCTGAGGAAAGAATATGAATCTCTTTTCACTTTAAAGGAA GATTC TGGGTTAGAGATTTCACCAAT | Double stranded DNA | CRISPR/Cas9 *S. cerevisiae*, repair template for 3xAID-3xHA-RSC2 | This study |
| 197bp_601DNA | ATCAGGTCGCTGTTCAATACATGCACAGGATGTATATATCTGA CACGTGCCTGGAGACTAGGGAGTAATCCCCTTGGCGGTTAAA ACGCGGGGGACAGCGCGTACGTGCGTTTAAGCGGTGCTAGA GCTGTCTACGACCAATTGAGCGGCCTCGGCACCGGGATTCTC CAGGGCGGCCGCGTATAGGGTCCATCGAT | Double stranded DNA | EMSA | Garcia-Saez et al., 2018 |
| 145bp_601DNA | ATCGATGTATATATCTGACACGTGCCTGGAGACTAGGGAGTAA TCCCCTTGGCGGTTAAAACGCGGGGGACAGCGCGTACGTGCG TTTAAGCGGTGCTAGAGCTGTCTACGACCAATTGAGCGGCCTC GGCACCGGGATTCTGAT | Double stranded DNA | EMSA | Garcia-Saez et al., 2018 |
| prJG1904_A-rich_F | GcatcagtcttataggAAGAAAAACAAAAAgtccatcgatctag | Oligonucleotide | EMSA | This study |
| prJG1905_A-rich_R | ctagatcgatggacTTTTTGTTTTTCTTcctataagactgatg | Oligonucleotide | EMSA | This study |
| prJG1906_TC-rich_F | GcatcagtcttataggTCTTCCTCTTCCTAgtccatcgatctag | Oligonucleotide | EMSA | This study |
| prJG1907_TC-rich_R | ctagatcgatggacTAGGAAGAGGAAGAcctataagactgatg | Oligonucleotide | EMSA | This study |

### Supplementary Table 5. BD-containing fungal proteins

| Specie | Protein | Name | UniProt ID |
|---|---|---|---|
| *Ascoidea rubescens* | Bdf1 | Bdf1_Arub | A0A1D2VIP6 |
| *Ascoidea rubescens* | Gcn5 | Gcn5_Arub | A0A1D2VKM0 |
| *Ascoidea rubescens* | Nto1 | Nto1_Arub | A0A1D2VQ20 |
| *Ascoidea rubescens* | Other | Other_Arub | A0A1D2VIG0 |
| *Ascoidea rubescens* | Rsc1 | Rsc1_Arub | A0A1D2VHR8 |
| *Ascoidea rubescens* | Rsc4 | Rsc4_Arub | A0A1D2VKQ1 |
| *Ascoidea rubescens* | Rsc58 | Rsc58_Arub | A0A1D2VNJ5 |
| *Ascoidea rubescens* | Snf2 | Snf2_Arub | A0A1D2VBJ0 |
| *Ascoidea rubescens* | Spt7 | Spt7_Arub | A0A1D2VMH6 |
| *Ascoidea rubescens* | Sth1 | Sth1_Arub | A0A1D2VHR6 |
| *Ascoidea rubescens* | Yta7 | Yta7_Arub | A0A1D2VPK3 |
| *Aspergillus fumigatus* | Bdf1 | Bdf1_Afum | Q4WPV0 |
| *Aspergillus fumigatus* | Gcn5 | Gcn5_Afum | Q4WQF6 |
| *Aspergillus fumigatus* | Nto1 | Nto1_Afum | Q4WGJ5 |
| *Aspergillus fumigatus* | Rsc1 | Rsc1_Afum | Q4WWE6 |
| *Aspergillus fumigatus* | Spt7 | Spt7_Afum | Q4WXX2 |
| *Aspergillus fumigatus* | Sth1 | Sth1_Afum | Q4WTW4 |
| *Aspergillus fumigatus* | Taf2 | Taf2_Afum | Q4X0S8 |
| *Aspergillus fumigatus* | Yta7 | Yta7_Afum | Q4WCI8 |
| *Candida albicans* | Bdf1 | Bdf1_Calb | Q5A4W8 |
| *Candida albicans* | Gcn5 | Gcn5_Calb | Q59PZ5 |
| *Candida albicans* | Nto1 | Nto1_Calb | Q5ANJ1 |
| *Candida albicans* | Other | Other_Calb | Q59R26 |
| *Candida albicans* | Rsc1 | Rsc1_Calb | A0A1D8PCV8 |
| *Candida albicans* | Rsc4 | Rsc4_Calb | A0A1D8PG88 |
| *Candida albicans* | Rsc58 | Rsc58_Calb | A0A1D8PG45 |
| *Candida albicans* | Snf2 | Snf2_Calb | A0A1D8PGK4 |
| *Candida albicans* | Spt7 | Spt7_Calb | A0A1D8PU33 |
| *Candida albicans* | Sth1 | Sth1_Calb | A0A1D8PJG4 |
| *Candida albicans* | Yta7 | Yta7_Calb | A0A1D8PP04 |
| *Candida arabinofermentans* | Bdf1 | Bdf1_Cara | A0A1E4STX8 |
| *Candida arabinofermentans* | Gcn5 | Gcn5_Cara | A0A1E4T6D2 |
| *Candida arabinofermentans* | Nto1 | Nto1_Cara | A0A1E4T2L6 |
| *Candida arabinofermentans* | Other | Other_Cara | A0A1E4SSS5 |
| *Candida arabinofermentans* | Rsc1 | Rsc1_Cara | A0A1E4SYH5 |
| *Candida arabinofermentans* | Rsc4 | Rsc4_Cara | A0A1E4SYN4 |
| *Candida arabinofermentans* | Rsc58 | Rsc58_Cara | A0A1E4T245 |
| *Candida arabinofermentans* | Snf2 | Snf2_Cara | A0A1E4SXM8 |
| *Candida arabinofermentans* | Spt7 | Spt7_Cara | A0A1E4SY86 |
| *Candida arabinofermentans* | Sth1 | Sth1_Cara | A0A1E4T627 |
| *Candida arabinofermentans* | Yta7 | Yta7_Cara | A0A1E4T0U9 |
| *Candida auris* | Bdf1 | Bdf1_Caur | A0A2H0ZCU0 |
| *Candida auris* | Gcn5 | Gcn5_Caur | A0A2H0ZCT4 |
| *Candida auris* | Nto1 | Nto1_Caur | A0A2H0ZNY6 |
| *Candida auris* | Other | Other_Caur | A0A2H0ZXE0 |
| *Candida auris* | Rsc1 | Rsc1_Caur | A0A2H0ZK45 |
| *Candida auris* | Rsc4 | Rsc4_Caur | A0A2H0ZM75 |
| *Candida auris* | Rsc58 | Rsc58_Caur | A0A2H0ZWR3 |
| *Candida auris* | Snf2 | Snf2_Caur | A0A2H1A183 |
| *Candida auris* | Spt7 | Spt7_Caur | A0A2H0ZKP1 |
| *Candida auris* | Sth1 | Sth1_Caur | A0A2H0ZVA0 |
| *Candida auris* | Yta7 | Yta7_Caur | A0A2H0ZK52 |
| *Candida glabrata* | Bdf1 | Bdf1_Cgla | Q6FWV7 |
| *Candida glabrata* | Gcn5 | Gcn5_Cgla | Q6FTW5 |
| *Candida glabrata* | Nto1 | Nto1_Cgla | Q6FMU0 |
| *Candida glabrata* | Rsc1 | Rsc1_Cgla | Q6FMX8 |
| *Candida glabrata* | Rsc2 | Rsc2_Cgla | Q6FXH7 |
| *Candida glabrata* | Rsc4 | Rsc4_Cgla | Q6FLM9 |
| *Candida glabrata* | Rsc58 | Rsc58_Cgla | Q6FY75 |
| *Candida glabrata* | Snf2 | Snf2_Cgla | Q6FJN8 |
| *Candida glabrata* | Spt7 | Spt7_Cgla | Q6FK24 |
| *Candida glabrata* | Sth1 | Sth1_Cgla | Q6FSQ1 |
| *Candida glabrata* | Yta7 | Yta7_Cgla | Q6FTQ6 |
| *Kluyveromyces lactis* | Bdf1 | Bdf1_Klac | Q6CWL8 |
| *Kluyveromyces lactis* | Gcn5 | Gcn5_Klac | Q6CXW4 |
| *Kluyveromyces lactis* | Nto1 | Nto1_Klac | Q6CP99 |
| *Kluyveromyces lactis* | Rsc1 | Rsc1_Klac | Q6CVB9 |
| *Kluyveromyces lactis* | Rsc4 | Rsc4_Klac | Q6CY60 |
| *Kluyveromyces lactis* | Rsc58 | Rsc58_Klac | Q6CJM7 |
| *Kluyveromyces lactis* | Snf2 | Snf2_Klac | Q6CVY8 |
| *Kluyveromyces lactis* | Spt7 | Spt7_Klac | Q6CNH2 |
| *Kluyveromyces lactis* | Sth1 | Sth1_Klac | Q6CLA5 |
| *Kluyveromyces lactis* | Yta7 | Yta7_Klac | Q6CXB0 |
| *Lachancea thermotolerans* | Bdf1 | Bdf1_Lthe | C5DFF4 |
| *Lachancea thermotolerans* | Gcn5 | Gcn5_Lthe | C5E1Y0 |
| *Lachancea thermotolerans* | Nto1 | Nto1_Lthe | C5DE16 |
| *Lachancea thermotolerans* | Rsc1 | Rsc1_Lthe | C5DBK3 |
| *Lachancea thermotolerans* | Rsc4 | Rsc4_Lthe | C5DGM3 |
| *Lachancea thermotolerans* | Rsc58 | Rsc58_Lthe | C5DI29 |

| | | | |
|---|---|---|---|
| *Lachancea thermotolerans* | Snf2 | Snf2_Lthe | C5DF84 |
| *Lachancea thermotolerans* | Spt7 | Spt7_Lthe | C5DIQ1 |
| *Lachancea thermotolerans* | Sth1 | Sth1_Lthe | C5DMI4 |
| *Lachancea thermotolerans* | Yta7 | Yta7_Lthe | C5DD84 |
| *Pneumocystis jirovecii* | Bdf1 | Bdf1_Pjir | A0A0W4ZCD1 |
| *Pneumocystis jirovecii* | Gcn5 | Gcn5_Pjir | A0A0W4ZUQ0 |
| *Pneumocystis jirovecii* | Nto1 | Nto1_Pjir | A0A0W4ZVR8 |
| *Pneumocystis jirovecii* | Other | Other_Pjir | A0A0W4ZQ45 |
| *Pneumocystis jirovecii* | Rsc1 | Rsc1_Pjir | A0A0W4ZFS0 |
| *Pneumocystis jirovecii* | Rsc4 | Rsc4_Pjir | A0A0W4ZET0 |
| *Pneumocystis jirovecii* | Spt7 | Spt7_Pjir | A0A0W4ZS82 |
| *Pneumocystis jirovecii* | Sth1 | Sth1_Pjir | A0A0W4ZRJ9 |
| *Saccharomyces cerevisiae* | Bdf1 | Bdf1_Scer | P35817 |
| *Saccharomyces cerevisiae* | Bdf2 | Bdf2_Scer | Q07442 |
| *Saccharomyces cerevisiae* | Gcn5 | Gcn5_Scer | Q03330 |
| *Saccharomyces cerevisiae* | Nto1 | Nto1_Scer | Q12311 |
| *Saccharomyces cerevisiae* | Rsc1 | Rsc1_Scer | P53236 |
| *Saccharomyces cerevisiae* | Rsc2 | Rsc2_Scer | Q06488 |
| *Saccharomyces cerevisiae* | Rsc4 | Rsc4_Scer | Q02206 |
| *Saccharomyces cerevisiae* | Rsc58 | Rsc58_Scer | Q07979 |
| *Saccharomyces cerevisiae* | Snf2 | Snf2_Scer | P22082 |
| *Saccharomyces cerevisiae* | Spt7 | Spt7_Scer | P35177 |
| *Saccharomyces cerevisiae* | Sth1 | Sth1_Scer | P32597 |
| *Saccharomyces cerevisiae* | Yta7 | Yta7_Scer | P40340 |
| *Scheffersomyces stipitis* | Bdf1 | Bdf1_Ssti | A3LZG2 |
| *Scheffersomyces stipitis* | Gcn5 | Gcn5_Ssti | A3LPA3 |
| *Scheffersomyces stipitis* | Nto1 | Nto1_Ssti | A3GHJ1 |
| *Scheffersomyces stipitis* | Other | Other_Ssti | A3LY64 |
| *Scheffersomyces stipitis* | Rsc1 | Rsc1_Ssti | A3LRA0 |
| *Scheffersomyces stipitis* | Rsc4 | Rsc4_Ssti | A3LMT3 |
| *Scheffersomyces stipitis* | Rsc58 | Rsc58_Ssti | A3LMV4 |
| *Scheffersomyces stipitis* | Snf2 | Snf2_Ssti | A3LTF0 |
| *Scheffersomyces stipitis* | Spt7 | Spt7_Ssti | A3GGX3 |
| *Scheffersomyces stipitis* | Sth1 | Sth1_Ssti | A3LZW6 |
| *Scheffersomyces stipitis* | Yta7 | Yta7_Ssti | A3LNS7 |
| *Schizosaccharomyces japonicus* | Bdf1 | Bdf1_Sjap | B6K2N3 |
| *Schizosaccharomyces japonicus* | Bdf2 | Bdf2_Sjap | B6K0Y3 |
| *Schizosaccharomyces japonicus* | Gcn5 | Gcn5_Sjap | B6K151 |
| *Schizosaccharomyces japonicus* | Nto1 | Nto1_Sjap | B6K0S0 |
| *Schizosaccharomyces japonicus* | Other | Other_Sjap | B6JXP6 |
| *Schizosaccharomyces japonicus* | Rsc1 | Rsc1_Sjap | B6K521 |
| *Schizosaccharomyces japonicus* | Rsc4 | Rsc4_Sjap | B6JVM5 |
| *Schizosaccharomyces japonicus* | Rsc58 | Rsc58_Sjap | B6K2V9 |
| *Schizosaccharomyces japonicus* | Snf2 | Snf2_Sjap | B6K540 |
| *Schizosaccharomyces japonicus* | Spt7 | Spt7_Sjap | B6K1J8 |
| *Schizosaccharomyces japonicus* | Sth1 | Sth1_Sjap | B6K7N8 |
| *Schizosaccharomyces pombe* | Bdf1 | Bdf1_Spom | Q9Y7N0 |
| *Schizosaccharomyces pombe* | Bdf2 | Bdf2_Spom | Q9HGP4 |
| *Schizosaccharomyces pombe* | Gcn5 | Gcn5_Spom | Q9UUK2 |
| *Schizosaccharomyces pombe* | Nto1 | Nto1_Spom | O74759 |
| *Schizosaccharomyces pombe* | Rsc1 | Rsc1_Spom | O74964 |
| *Schizosaccharomyces pombe* | Rsc4 | Rsc4_Spom | Q09948 |
| *Schizosaccharomyces pombe* | Rsc58 | Rsc58_Spom | Q10412 |
| *Schizosaccharomyces pombe* | Snf2 | Snf2_Spom | O94421 |
| *Schizosaccharomyces pombe* | Spt7 | Spt7_Spom | P87152 |
| *Schizosaccharomyces pombe* | Sth1 | Sth1_Spom | Q9UTN6 |
| *Wickerhamomyces ciferrii* | Bdf1 | Bdf1_Wcif | K0KYG4 |
| *Wickerhamomyces ciferrii* | Gcn5 | Gcn5_Wcif | K0KPB8 |
| *Wickerhamomyces ciferrii* | Nto1 | Nto1_Wcif | K0KW43 |
| *Wickerhamomyces ciferrii* | Other | Other_Wcif | K0KBT4 |
| *Wickerhamomyces ciferrii* | Rsc1 | Rsc1_Wcif | K0KL11 |
| *Wickerhamomyces ciferrii* | Rsc4 | Rsc4_Wcif | K0KL87 |
| *Wickerhamomyces ciferrii* | Rsc58 | Rsc58_Wcif | K0KVS3 |
| *Wickerhamomyces ciferrii* | Snf2 | Snf2_Wcif | K0KQW9 |
| *Wickerhamomyces ciferrii* | Spt7 | Spt7_Wcif | K0KIS0 |
| *Wickerhamomyces ciferrii* | Sth1 | Sth1_Wcif | K0KK36 |
| *Wickerhamomyces ciferrii* | Yta7 | Yta7_Wcif | K0KJH7 |
| *Yarrowia lipolytica* | Bdf1 | Bdf1_Ylip | Q6C571 |
| *Yarrowia lipolytica* | Gcn5 | Gcn5_Ylip | Q8WZM0 |
| *Yarrowia lipolytica* | Nto1 | Nto1_Ylip | Q6C7S8 |
| *Yarrowia lipolytica* | Other | Other_Ylip | Q6C6Z3 |
| *Yarrowia lipolytica* | Rsc1 | Rsc1_Ylip | Q6C8C8 |
| *Yarrowia lipolytica* | Rsc4 | Rsc4_Ylip | Q6C3T3 |
| *Yarrowia lipolytica* | Rsc58 | Rsc58_Ylip | Q6C7M7 |
| *Yarrowia lipolytica* | Snf2 | Snf2_Ylip | Q6C828 |
| *Yarrowia lipolytica* | Spt7 | Spt7_Ylip | Q6C3U2 |
| *Yarrowia lipolytica* | Sth1 | Sth1_Ylip | Q6CDE1 |

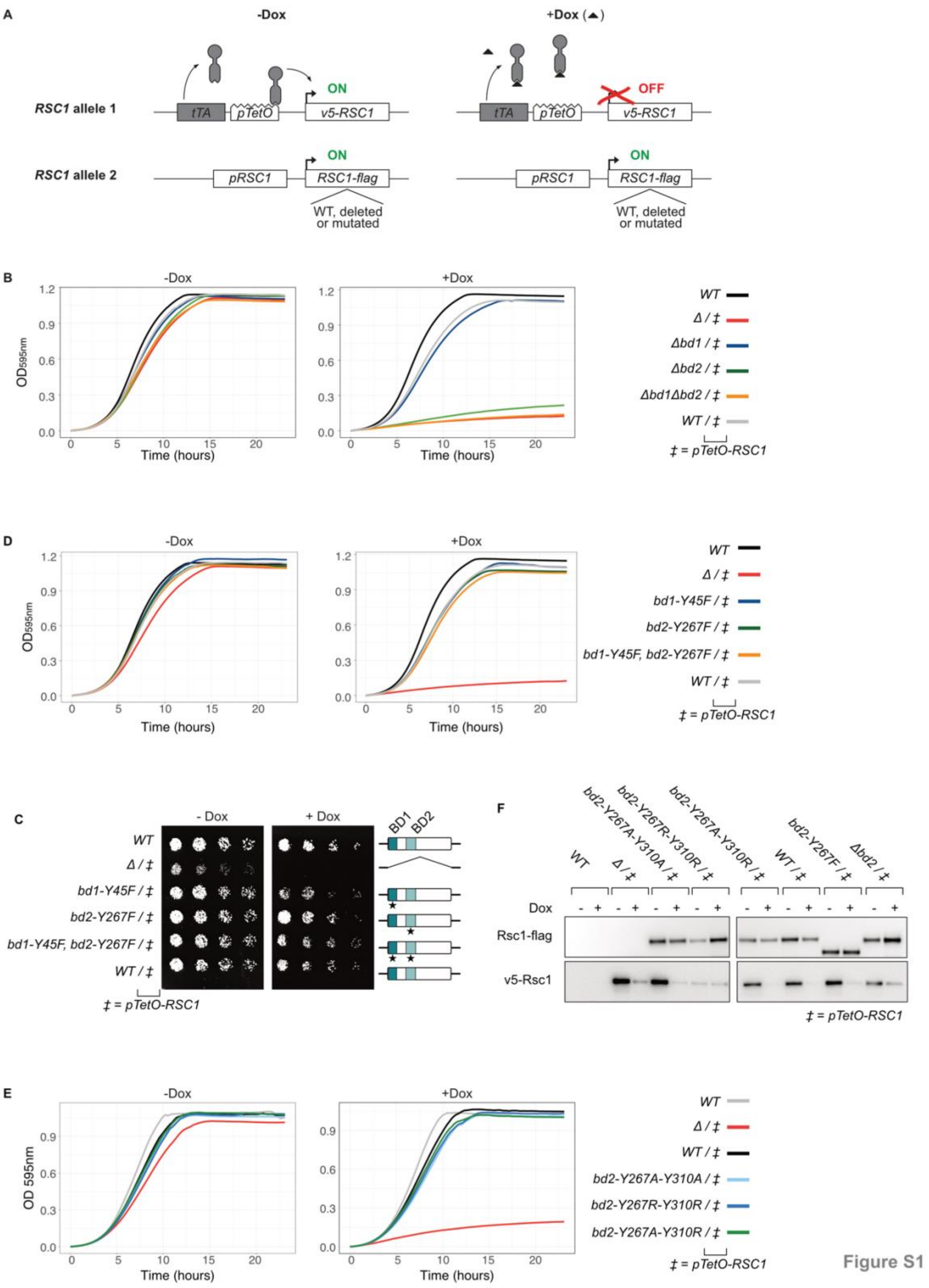


**Figure S1.**

**A.** *C. albicans RSC1* conditional alteration using the Tet-Off system. One *RSC1* allele is placed under the *pTetO* promoter. In the absence of doxycycline (Dox), *pTetO* is activated by a recombinant transcription factor (tTA) and allele 1 is expressed. Addition of exogenous Dox prevents tTA from binding to *pTetO*, resulting in transcriptional repression of allele 1. *RSC1* from allele 2 is controlled by its endogenous promoter, the ORF has been modified to express either a WT or mutated Rsc1 variant, or has been fully deleted. Rsc1 expressed from each allele has been fused to an epitope tag (v5 or flag) to allow western-blot detection.

**B.** Growth assays performed with the same *C. albicans* strains as in **Figure 1A**. Liquid cultures where grown -/+ Dox, the optical density ($OD_{595nm}$) was monitored over time. Biological duplicates have been averaged.

**C.** Growth assays as in panel A, performed with the same *C. albicans* strains as in **Figure 1B**.

**D.** Western-blot performed with the same strains as in **Figure 1B**. The *pTetO-RSC1* allele expresses v5-Rsc1 protein. The other allele expresses Rsc1-flag, WT or mutated.

**E.** *C. albicans* Colony formation assay upon conditional alteration of *RSC1*, performed in duplicate, as in **Figure 1A**.

**F.** Growth assays performed with the same *C. albicans* strains as in panel D.

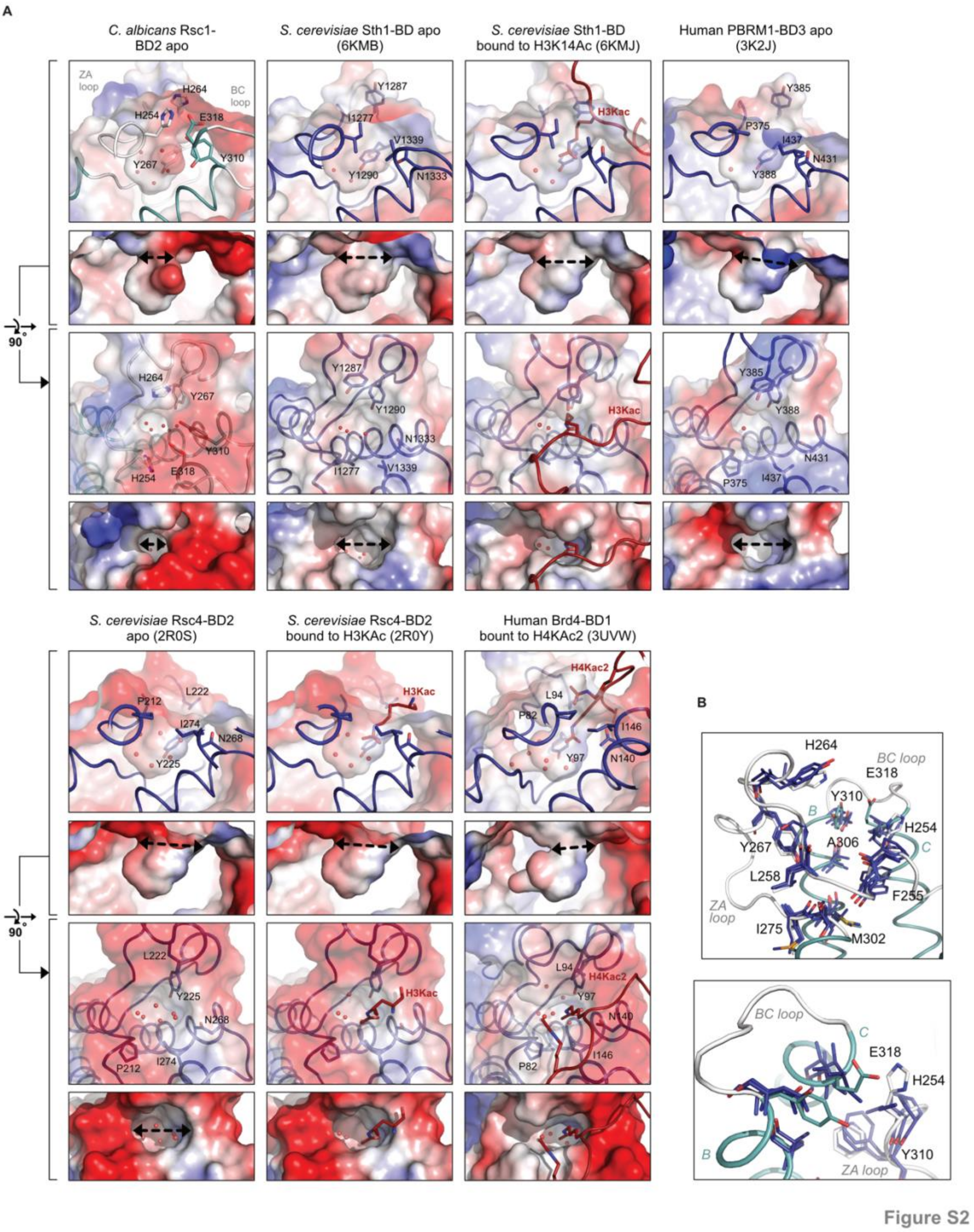


**Figure S2.**
**A.** Detailed view of Rsc1-BD2 putative binding pocket compared with other BDs in their apo or peptide-bound form. For each BD, the PDB code is indicated (23,25,98).
**B.** Rsc1-BD2 is aligned to Sth1-BD, PBRM1-BD3, Rsc4-BD2 in their apo forms. Rsc1-BD2 is coloured in light teal for helices and white for loops, residues involved in the pocket architecture are shown as sticks. Other BDs are all coloured in dark blue with only pocket residues shown as sticks.

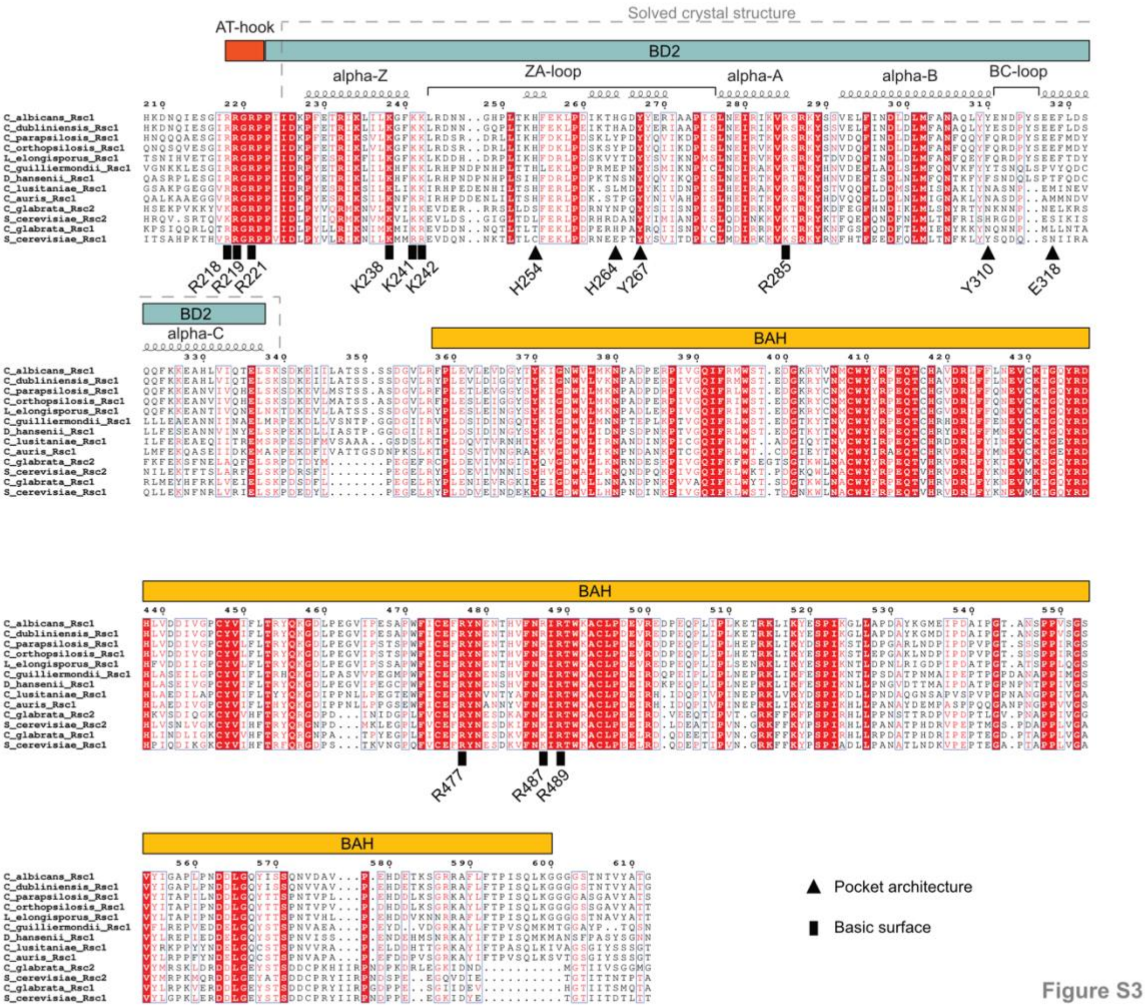


**Figure S3**.

Sequence alignment of Rsc1 homologues from multiple yeast species, showing the AT-hook, BD2 and BAH domains. Residues are numbered according to *C. albicans* Rsc1. The secondary structure of BD2 is annotated according to the *C. albicans* Rsc1 crystal structure determined in this study. Fully conserved positions are highlighted in red, positions with conserved properties are with red letters. Triangles indicate residues determining BD2 pocket architecture, rectangles indicate residues responsible for the basic surface.

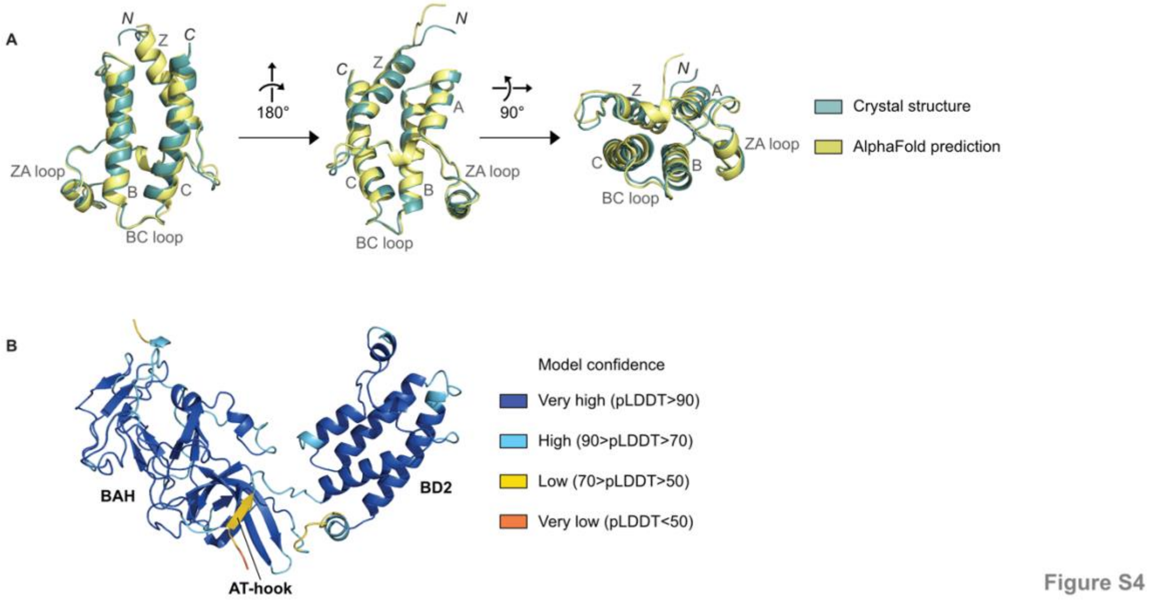


**Figure S4.**

**A.** 3D alignment of *C. albicans* Rsc1-BD2 structures, experimentally determined using X-ray crystallography (light teal) and predicted by AlphaFold (yellow). Protein backbones aligned with and RMSD of 0.680 Å for 105 C$\alpha$.

**B.** 3D model of *C. albicans* Rsc1 213-599 predicted by the AlphaFold database, coloured by model confidence.

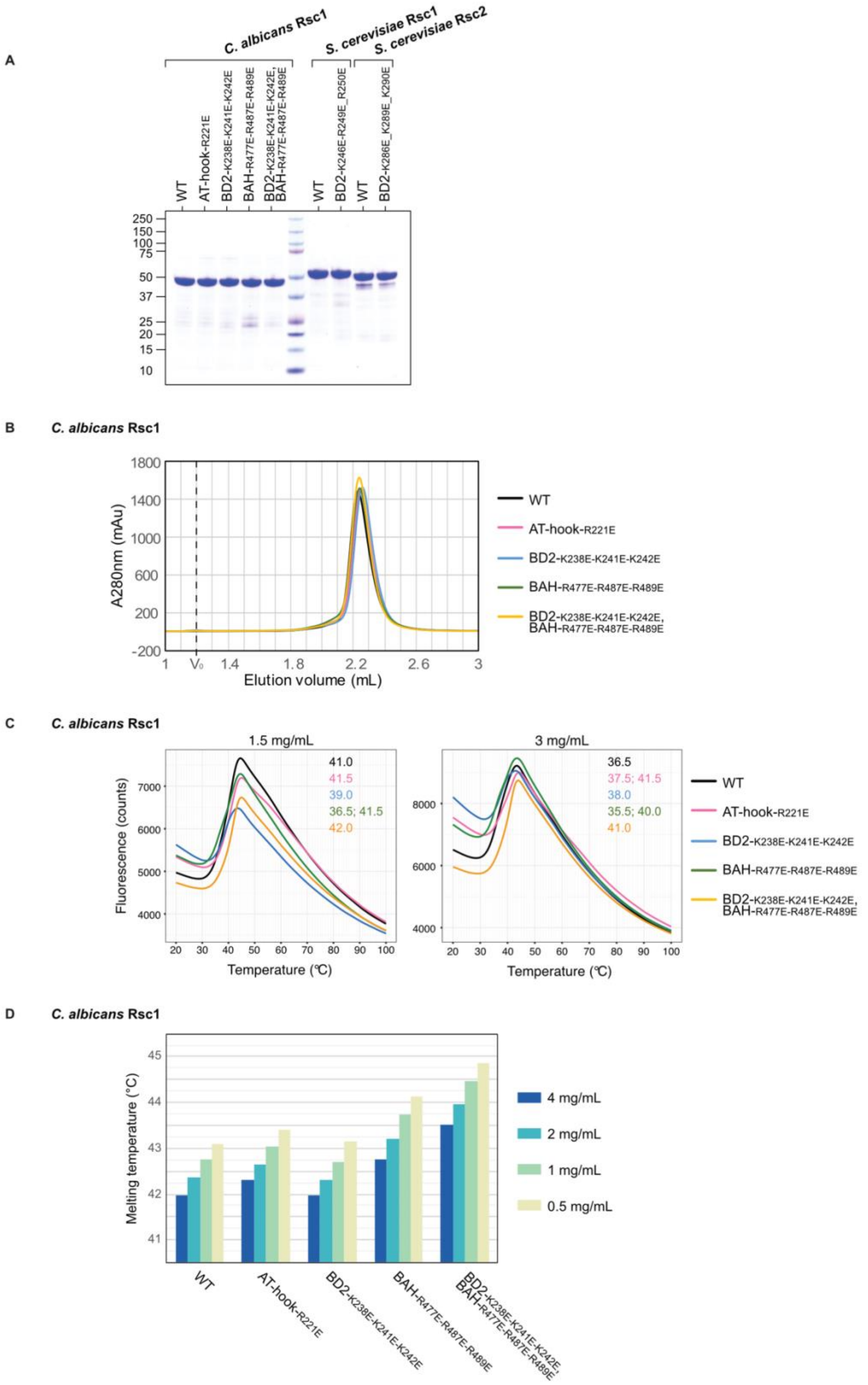


Figure S5

**Figure S5**.

**A.** SDS-PAGE analysis of purified recombinant proteins produced in *E. coli* and used for EMSA (Figure 2D and 5C-E). 5 µg of protein were loaded on the gel.

**B.** Gel filtration analysis of *C. albicans* Rsc1 recombinant proteins. 250 µg of WT or mutated proteins eluted with similar profiles from a Superdex 200 Increase 5/150 GL. $V_0$ indicates the void volume of the column.

**C.** Thermal Shift Assay of *C. albicans* Rsc1 recombinant proteins at 1.5 and 3 mg/mL. The melting temperatures (°C) obtained from each curve are indicated on each plot.
**D.** Nano Differential Scanning Fluorimetry analysis of *C. albicans* Rsc1 recombinant proteins.

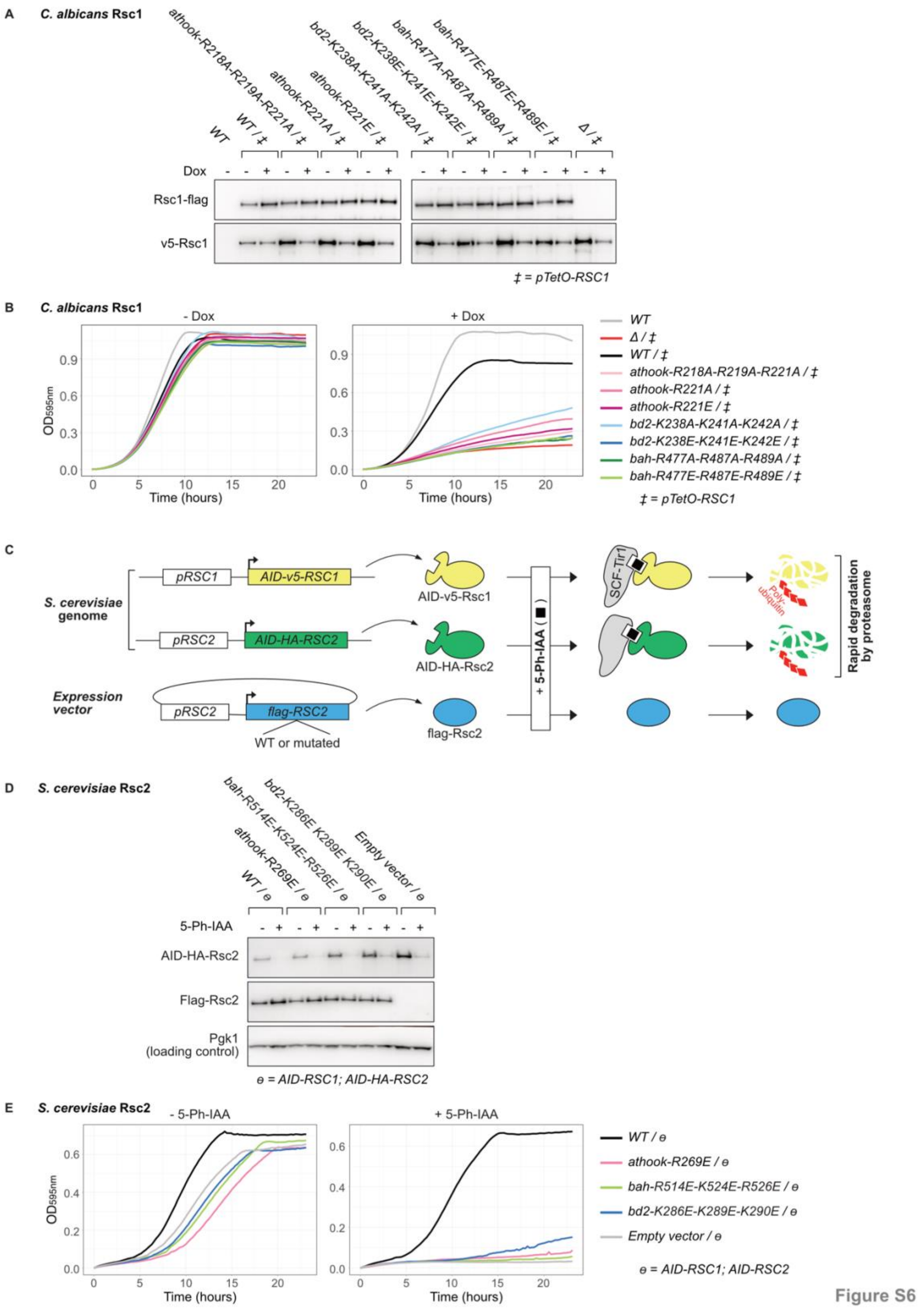


**Figure S6.**

**B.** Western-blot performed with the same *C. albicans* strains as in Figure 2F. The *pTetO-RSC1* allele expresses v5-Rsc1 protein. The other allele expresses Rsc1-flag, WT or mutated.

**A.** Growth assays performed with the same *C. albicans* strains as in **Figure 2F**. Liquid cultures where grown ± Dox, $OD_{595nm}$ was monitored over time. Biological duplicates have been averaged.
**C.** *S. cerevisiae RSC1* and *RSC2* conditional alteration using a custom Auxin Induced Degron (AID) system. An AID domain has been fused to each *RSC1* and *RSC2* paralogue, at its endogenous locus, using marker-less CRISPR/Cas9. A copy of *RSC2* is also expressed from a vector (*pRS316*), in its WT or mutated from. Each strain also carries a gene encoding $OsTir1F_{74A}$ (Tir1). Upon addition of 5-Ph-IAA, an auxin analogue, Tir1 mediates interaction of the AID domain, with the SCF complex, leading to poly-ubiquitination and degradation by the proteasome.
**E.** Western-blot performed with the same *S. cerevisiae* strains as in **Figure 2G**. Rsc2 expressed from its endogenous locus is fused to an AID-HA tag; Flag-Rsc2, WT or mutated, is expressed from a plasmid.
**D.** Growth assays performed with the same *S. cerevisiae* strains as in **Figure 2G**. Liquid cultures where grown ± 5Ph-IAA, $OD_{595nm}$ was monitored over time. Biological duplicates have been averaged.

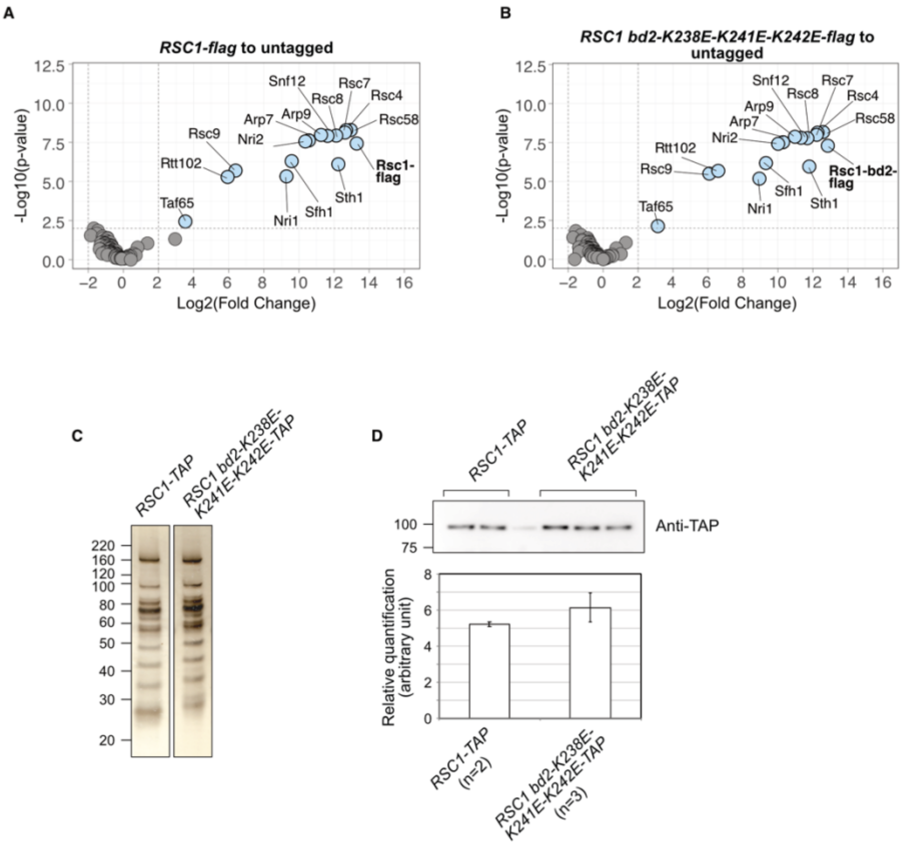


Figure S7

**Figure S7.**

**A-B.** Identification of proteins co-immunoprecipitated with Rsc1-flag (**A**) or Rsc1-bd2-$_{K238E-K241E-K242E}$-flag (**B**) compared to the untagged form, by mass spectrometry-based label-free quantitative proteomics. Analysis from triplicate experiments is represented as volcano plots; dashed lines show cutoffs at -Log10(P- value)=2 and Log2(Fold change)=2 ; proteins above these thresholds are coloured in blue (P-value>0.01, Fold change<4).

**C.** Silver staining detection of proteins eluted after TAP purification of the endogenous RSC complex from *C. albicans* strains expressing Rsc1-TAP or Rsc1-bd2-$_{K238E-K241E-K242E}$-TAP.

**D.** Quantification of Rsc1-TAP and Rsc1-bd2-$_{K238E-K241E-K242E}$-TAP in proteins eluted after TAP purification, by western-blot analysis. Rsc1 proteins are detected using an anti-TAP tag antibody (top panel) and the signal was quantified (bottom panel).

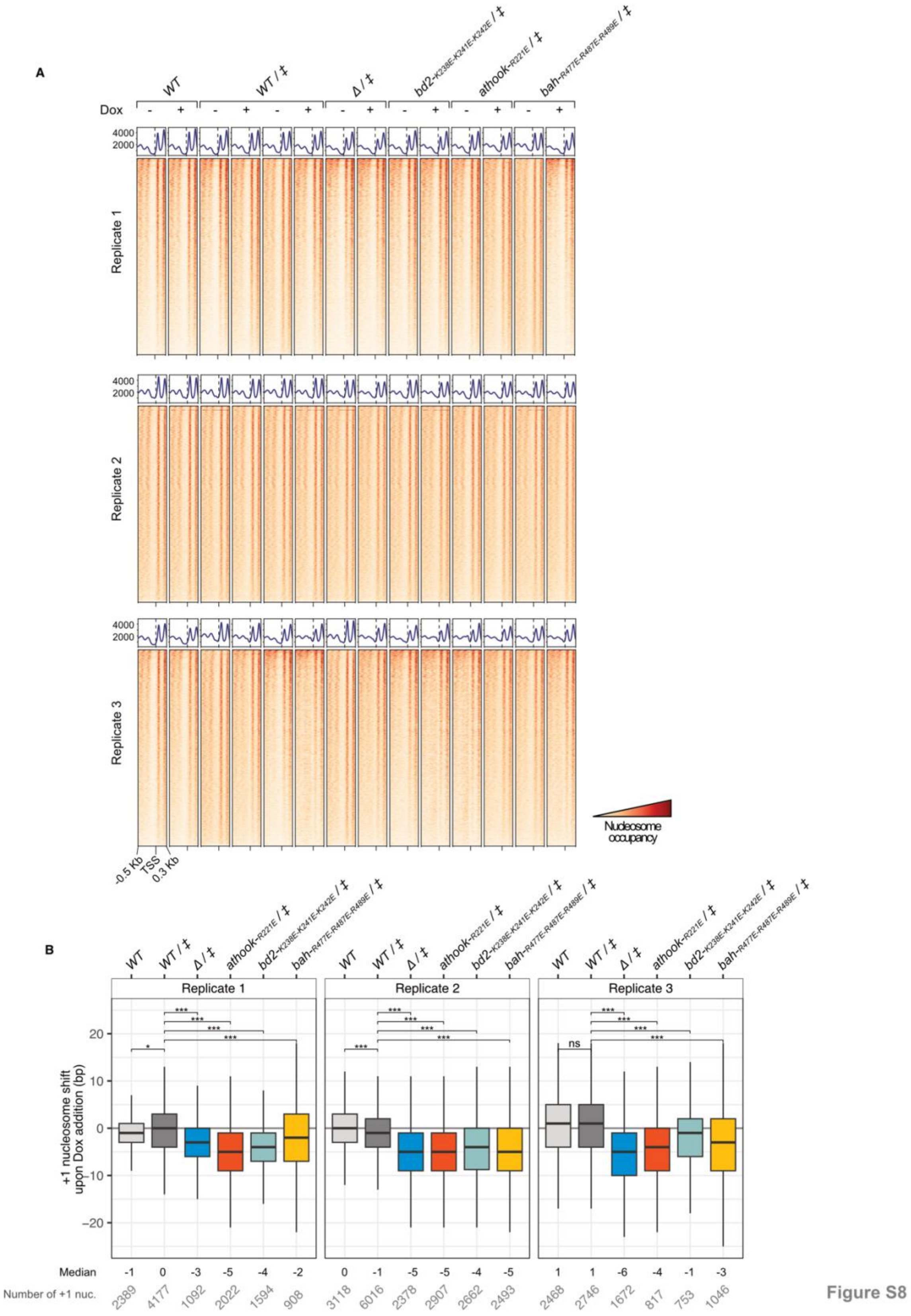


**Figure S8.**

**A.** Nucleosomal occupancy averaged profile (top) and heatmap (bottom) of each MNase-seq samples, over all protein coding genes (6221 genes). Genes are aligned to TSS with -0.3 and 0.2 Kb flanks.

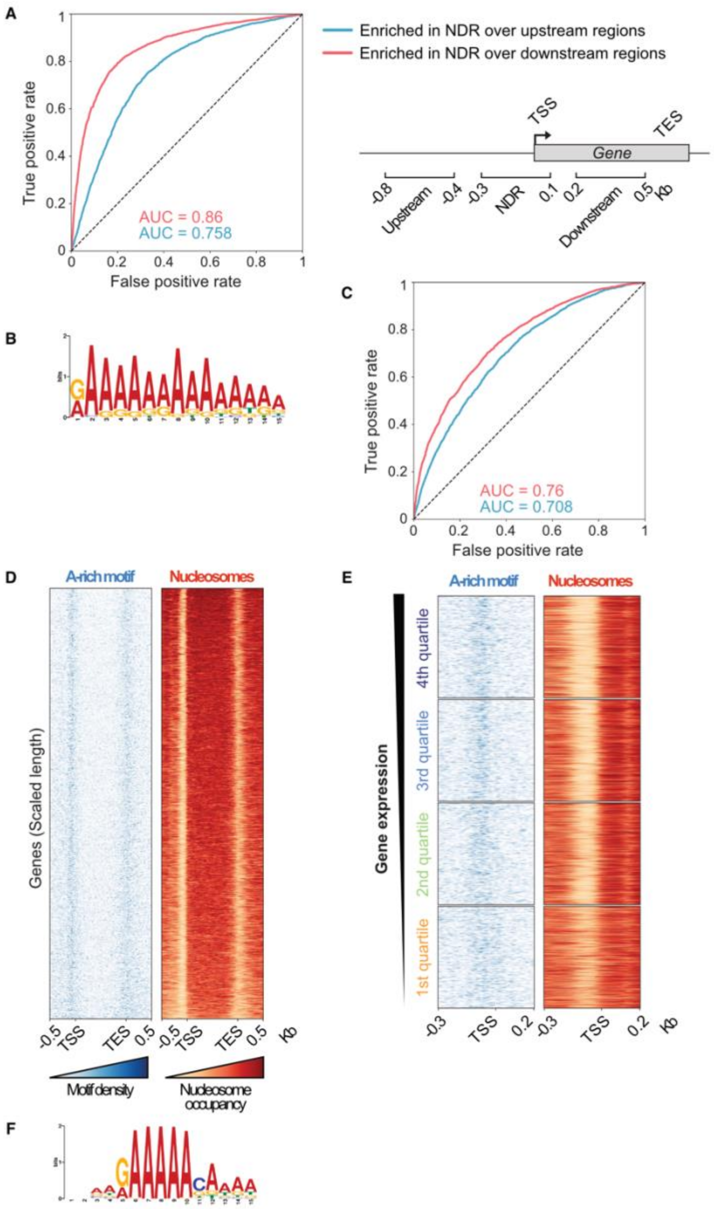


Figure S9

**Figure S9.**

**A.** Receiver operating characteristic (ROC) curve for the *de novo* polyA motif enriched in *C. albicans* NDR (Figure 4A), showing enrichment of the motif in the NDR compared to adjacent genomic regions. Intervals used for NDR and adjacent regions are illustrated on the right. The values of the Area Under the Curve (AUC) are indicated.

**B.** A-rich Motif identified *de novo* in the NDR region (-300 to +100 bp around the TSS) of *S. cerevisiae* protein coding genes.

**C.** As in panel **A**, for the *de novo* polyA motif enriched in *S. cerevisiae* NDR.

**D.** Heatmaps of the A-rich motif density (left side, coloured blue) and nucleosomal occupancy (right side, coloured orange) over all protein coding genes in *S. cerevisiae*. Gene length is scaled from TSS to TES, with 0.5 Kb flanks. The nucleosomal occupancy is from (93).

**E.** Average profiles of motif density (blue curve, number of motifs) and nucleosomal occupancy (orange curve, RPKM) over *S. cerevisiae* genes sorted and stratified by gene expression, based on (94).

**F.** Motif obtained by ampDAP-seq using *in vitro* produced *S. cerevisiae* Rsc2 protein and exogenous PCR-amplified genomic DNA.
